# HLA-C-restricted presentation of a conserved bacterial epitope to an innate NK cell receptor

**DOI:** 10.1101/550889

**Authors:** Malcolm J. W. Sim, Sumati Rajagopalan, Daniel M. Altmann, Rosemary J. Boyton, Peter D. Sun, Eric O. Long

## Abstract

The killer-cell Ig-like receptor (KIR) family, expressed mainly in natural killer (NK) cells, includes an activation receptor of unknown function, KIR2DS4. Here we show that KIR2DS4 is restricted by HLA-C*05:01 with a strong preference for tryptophan at position 8 of 9-mer peptides. ‘Self’ peptides with Trp8 eluted from HLA-C*05:01 are rare and only one out of 12 bound KIR2DS4. An HLA-C*05:01-peptide complex that bound KIR2DS4 was sufficient for strong activation of primary KIR2DS4^+^ NK cells, independently of coactivation by other receptors and of prior NK cell licensing. A highly conserved sequence in bacterial recombinase A, which is essential for DNA repair and survival, includes an epitope that bound to HLA-C*05:01 and activated KIR2DS4^+^ NK cells. Thus, in addition to their established role in defense against viruses and cancer, NK cells may have also evolved to detect and respond to hundreds of bacterial species through recognition of a conserved RecA epitope.

## Introduction

MHC class I molecules (MHC-I) play critical roles in innate and adaptive immunity. MHC-I molecules present short peptides, commonly 8-11 amino acids in length, which are surveilled by alpha-beta T cell receptors expressed by CD8 T cells. MHC-I also serves as a critical regulator of natural killer (NK) cells, innate immune cytotoxic cells with the capacity to produce pro-inflammatory cytokines (Long et al., 2013; Vivier et al., 2011). Following the ‘missing self’ hypothesis, MHC-I binding inhibitory receptors expressed by NK cells detect loss of MHC-I leading to NK cell activation (Ljunggren and Karre, 1990). Additionally, interactions between inhibitory receptors and MHC-I dictate the effector potential of NK cells via a process known as ‘education’ or ‘licensing’ (Elliott and Yokoyama, 2011; Long et al., 2013). NK cells have established roles in immune defense against cancers and viral infections, where loss or down-regulation of MHC-I is common (Morvan and Lanier, 2016; Vidal et al., 2011). The functions of MHC-I binding NK cell inhibitory receptors appear conserved across species and different families of receptors.

In humans, the major NK receptors for HLA-I (human MHC-I) are CD94:NKG2A, which binds HLA-E, and the killer-cell immunoglobulin-like receptors (KIR). There are 14 KIR genes which encode activating and inhibitory receptors. The ligands for inhibitory KIR are well defined as groups of HLA-A, -B or -C allotypes, each with a common epitope. All HLA-C allotypes carry either the C1 or C2 epitopes, which are ligands for the inhibitory receptors KIR2DL2/3 and KIR2DL1 respectively (Parham, 2005). KIR bind the peptide exposed face of HLA-I towards the C-terminal end of the peptide, incorporating peptide into the binding site and all HLA-C binding KIR studied to date demonstrate a degree of peptide selectivity (Boyington et al., 2000; Boyington and Sun, 2002; Fan et al., 2001; Rajagopalan and Long, 1997; Sim et al., 2017; Stewart et al., 2005). In contrast to the inhibitory KIR, definitive functional ligands for activating KIR are still lacking.

The KIR genes are organized into two broad haplotypes, KIR A and KIR B, which differ by gene content. The simpler KIR A haplotype contains only one activating receptor *KIR2DS4*, while the B haplotype is characterized by variable gene content including multiple activating receptor genes (Parham, 2005). The two haplotypes have a similar worldwide frequency, but differ between populations, such that KIR A homozygotes are not rare and for whom *KIR2DS4* is the only activating KIR they carry. Due to variability of KIR haplotypes and the fact that HLA-I and KIR are on different chromosomes, individuals can express orphan receptors or express ligands without the corresponding KIR. Consequently, gene association studies have linked the presence or absence or KIR–ligand pairs with many disease processes including viral infections, autoimmunity and cancer (Bashirova et al., 2011; Boyton and Altmann, 2007; Hiby et al., 2004; Khakoo et al., 2004; Parham, 2005; Parham and Moffett, 2013). Additionally, activating KIR with the ability to bind HLA-C appear to have a protective role against disorders of pregnancy (Hiby et al., 2004; Kennedy et al., 2016; Nakimuli et al., 2015).

The *KIR2DS4* locus is not fixed and two major alleles exist that encode either the full-length receptor (KIR2DS4-fl) or a version with a deletion (KIR2DS4-del). KIR2DS4-del encodes a 22-bp deletion leading to an early stop codon creating a truncated soluble protein with no recorded HLA-I binding (Graef et al., 2009; Maxwell et al., 2002). KIR2DS4-fl is an HLA-I binding receptor and binds a subset of C1 and C2 HLA-C allotypes, in contrast to other KIR2D receptors, which dominantly bind C1 or C2 (Graef et al., 2009). This previous report identified KIR2DS4 ligands via a binding assay using soluble KIR molecules and many HLA-A, -B and -C proteins bound to beads (Hilton et al., 2015a). This method has proved useful to screen many HLA-I allotypes at once, but the sequence and diversity of peptides presented on the beads is unknown. Furthermore, it is not clear whether HLA-C constitutes a functional ligand for KIR2DS4 or how peptide sequence contributes to KIR2DS4 binding. Indeed, the only known functional ligand for KIR2DS4 is HLA-A*11:02 (Graef et al., 2009).

Carrying KIR2DS4-fl is associated with protection from pre-eclampsia and glioblastoma, and with higher viral loads and faster progression to AIDS, in HIV infection (Dominguez-Valentin et al., 2016; Kennedy et al., 2016; Merino et al., 2014; Olvera et al., 2015). There is a clear need to define functional ligands for KIR2DS4 to fully understand its role in these disease processes and in the regulation of NK cells more generally. HLA-C*05:01, a C2 allotype, was reported to bind KIR2DS4-fl (Graef et al., 2009). Henceforth, we refer to KIR2DS4-fl as KIR2DS4 unless in direct comparison to KIR2DS4-del. The aim of this study was to establish whether HLA-C*05:01 is a functional ligand for KIR2DS4 and if peptide sequence influences KIR2DS4 binding. We find that KIR2DS4 binds HLA-C*05:01 in a highly peptide selective manner and that this binding potently activates KIR2DS4^+^ NK cells. Further, we link peptide specific recognition of HLA-C*05:01 by KIR2DS4 to a protein highly conserved among bacteria pathogenic in humans.

## Results

### Highly selective binding of KIR2DS4 to HLA-C*05:01 loaded with a 9mer peptide with tryptophan at position 8

Binding of a soluble KIR2DS4–IgG1-Fc fusion protein (KIR2DS4-Fc) was tested with the HLA-I deficient cell line 721.221 (221) transfected with HLA-C*05:01 (221–C*05:01) or HLA-C*04:01 (221–C*04:01; SFig. 1A). While KIR2DL1-Fc bound both cell lines strongly, no binding to KIR2DS4-Fc was detected. KIR2DL1 bound HLA-C*05:01 in the context of many different peptide sequences (Sim et al., 2017). To test whether KIR2DS4 binding may be more peptide specific, we examined KIR2DS4-Fc binding to HLA-C*05:01 loaded with individual peptide sequences, using a TAP–deficient cell line expressing HLA-C*05:01, as described (Sim et al., 2017). We tested 46 ‘self’ peptides previously eluted from purified HLA-C*05:01, none conferred binding of KIR2DS4-Fc to HLA-C*05:01 (SFig. 1B,C). However, ‘self’ peptide P2 (IIDKSGSTV) substituted with Ala and Tyr at positions p7 and p8 (P2-AY) conferred strong KIR2DS4-Fc binding to HLA-C*05:01 (SFig. 1B,C). P2-AY was synthesized as part of screen to test the impact of peptide sequence at p7 and p8 on inhibitory KIR binding (Sim et al., 2017). Nine additional amino acid substitutions at p7 and p8 of peptide P2 failed to bind to KIR2DS4, which suggested a contribution of the p8 Tyr. HLA-C*08:02 is a C1 allotype that differs from HLA-C*05:01 by only the C1 and C2 epitope (amino acids at positions 77 and 80) and is stabilized by peptides eluted from HLA-C*05:01 (Sim et al., 2017). However, KIR2DS4 did not bind HLA-C*08:02 when loaded with P2-AY, suggesting the C2 epitope positively contributed to KIR2DS4 binding to HLA-C*05:01 (SFig. 1D). As Tyr is a large aromatic residue, we tested two other large aromatic residues Phe (P2-AF) and Trp (P2-AW) at p8 and used Val as a control (P2-AV; Fig. 1A). All four peptides stabilized HLA-C*05:01 on 221-C*05:01-ICP47 cells to a similar extent and conferred strong KIR2DL1-Fc binding (Fig. 1A,B). Substitution of p8 Tyr to Phe reduced KIR2DS4-Fc binding, while substitution to Trp substantially increased binding of KIR2DS4-Fc (Fig. 1C,D). To confirm this interaction, soluble HLA-C*05:01 that was refolded with peptides P2-AV or P2-AW was produced as tetramers. Tetramers bound similarly to 293T cells transiently transfected with KIR2DL1 (SFig. 1F). In contrast, only HLA-C*05:01 tetramers refolded with P2-AW bound cells transfected with KIR2DS4 and DAP12 (Fig. 1E,F). Negligible binding of HLA-C*05:01 tetramers occurred with cells transfected with DAP12 alone. Collectively, these data show that KIR2DS4 binds HLA-C*05:01 with a high peptide selectivity. Out of 61 different peptide sequences tested, only P2-AF, P2-AY and P2-AW conferred measurable KIR2DS4 binding to HLA-C*05:01.

**Figure. 1.**
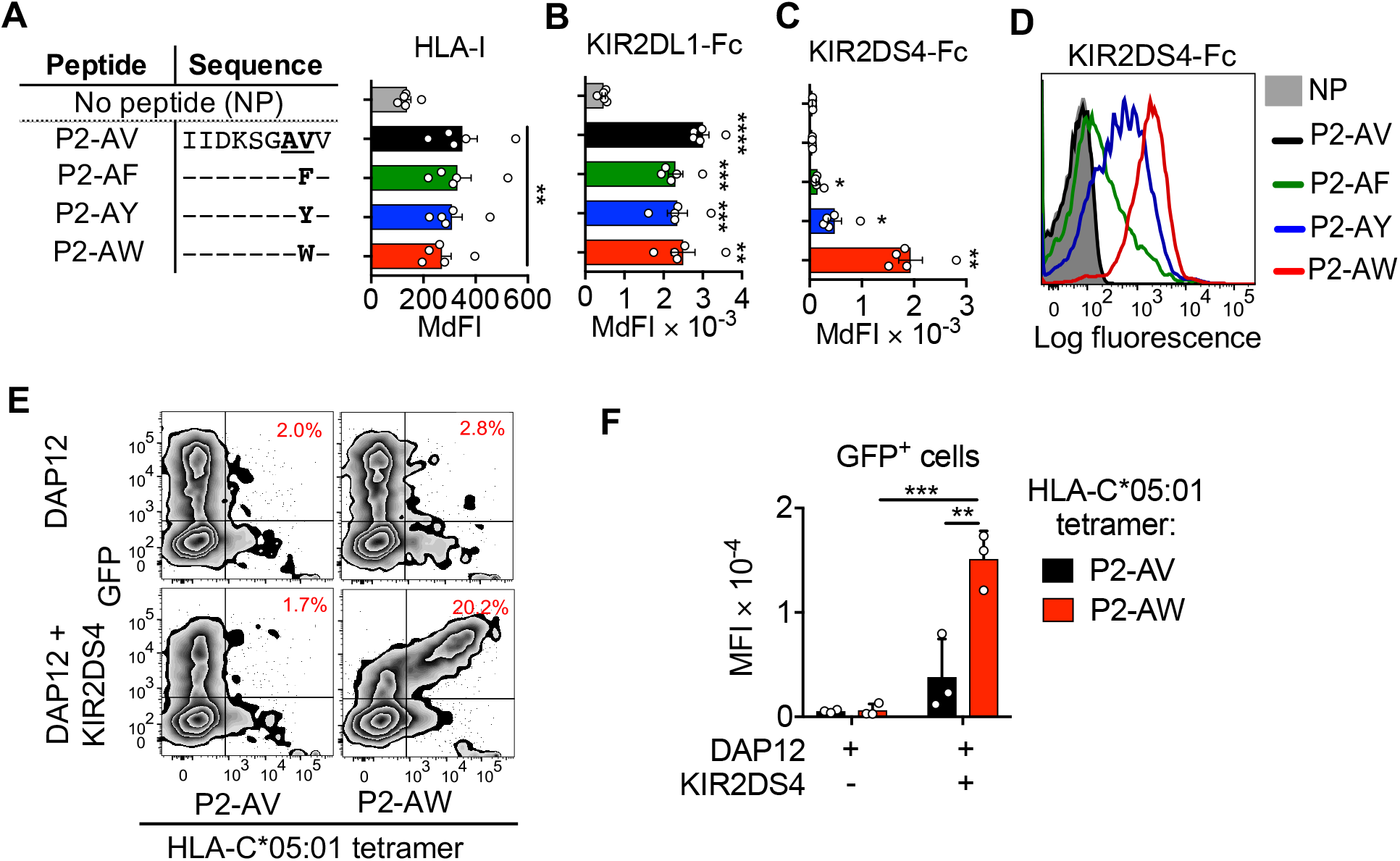
Binding of KIR2DS4 to HLA-C*05:01 loaded with a peptide with tryptophan at position 8. **(A)** Stabilization of HLA-I expression on 221–C*05:01-ICP47 cells incubated overnight at 26°C with 100 μM of peptides P2-AV, P2-AF, P2-AY and P2-AW. Amino acids identical to the P2-AV sequence are indicated with –. Data shown as median fluorescence intensity (MdFI), n=5. **(B)** KIR2DL1-Fc and **(C)** KIR2DS4-Fc binding to 221–C*05:01-ICP47 cells incubated with peptides as shown in **(B)** or NP, n=5. **(D)** Flow cytometry histograms displaying KIR2DS4-Fc binding to 221–C*05:01-ICP47 cells from a representative experiment, from a total of 5 independent experiments. **(E)** Flow cytometry bi-plots displaying HLA-C*05:01-tetramer binding to 293T cells transfected with separate vectors (pIRES2-eGFP) containing cDNA encoding DAP12 and KIR2DS4, or DAP12 only. HLA-C*05:01 tetramers were refolded with P2-AV or P2-AW. Data are from one representative experiment out of 3 independent experiments. **(F)** HLA-C*05:01-tetramer binding to 293T cells as in **(E)**, displayed as mean fluorescence intensity (MFI) from three independent experiments. Statistical significance was assessed by one-way ANOVA with Tukey’s multiple comparisons test (A-C) and two-way **(F)** ANOVA with Dunnett’s multiple comparisons test, *, P < 0,05, **, P < 0.01, ***, P < 0.001, ****, P < 0.0001. In A-C, significance is indicated in comparison to NP.

### Peptide–specific activation of KIR2DS4^+^ NK cells by HLA-C*05:01

We next tested the capacity of peptide:HLA-C*05:01 complexes to activate KIR2DS4^+^ NK cells in degranulation assays. Primary resting NK cells from KIR2DS4^+^ donors were mixed with 221-C*05:01-ICP47 cells loaded with peptides P2-AV, P2-AF, P2-AY or P2-AW. To eliminate the contribution KIR2DL1/S1 binding to HLA-C*05:01, NK cells were gated as CD56^dim^ and KIR2DL1/S1 (EB6) negative. KIR2DS4^+^, but not KIR2DS4^−^ NK cells displayed increased degranulation in response to cells loaded with P2-AY and P2-AW (SFig. 2A,B).

Due to the variegated expression of inhibitory NKG2A and KIR, KIR2DS4^+^ and KIR2DS4^−^ NK cells include both licensed and unlicensed NK cells. Licensing endows NK cell subsets with greater capacity to degranulate in the absence of MHC-I. However, activation of KIR2DS4^+^ NK cells requires expression of HLA-C, thus it is not clear how NK cell licensing may impact activation of KIR2DS4^+^ NK cells. To examine the role of NK cell licensing on activation of KIR2DS4^+^ NK cells, we gated CD56^dim^ KIR2DL1/S1^−^ NK cells into four subsets based on the expression of KIR2DS4 (S4) and other MHC-I binding receptors (KIR2DL2/L3/S2, NKG2A and KIR3DL1/S1), termed receptor positive (R^+^) and receptor negative (R^−^) (Fig. 2A). All donors carry the ligand for NKG2A (HLA-E), while donors vary for the presence of the ligands for KIR3DL1 and KIR2DL2/3. Thus, R^+^ NK cells are expected to be licensed and degranulate well in response 221-C*05:01-ICP47 cells as they lack ligands for KIR2DL2/3, NKG2A and KIR3DL1. R^−^ NK cells lack all licensing receptors and are expected to be degranulate weakly in response 221-C*05:01-ICP47 cells. As expected, licensed R^+^ NK cells degranulated well in response to 221-C*05:01-ICP47 cells, while the R^−^ NK cells degranulated poorly irrespective of KIR2DS4 expression (Fig. 2B,C). In contrast, KIR2DS4^+^ NK cells degranulated strongly in response to cells loaded with P2-AW (Fig. 2B). As expected, cells loaded with P2-AV did not stimulate KIR2DS4^+^ NK cells, while P2-AF and P2-AY did stimulate KIR2DS4^+^ NK cells to a lesser extent than P2-AW (Fig. 2B,C). Remarkably, the potent activation of KIR2DS4^+^ NK cells in response to cells loaded with P2-AW was similar between R^+^ and R^−^ subsets. Thus, even unlicensed R^−^ NK cells can respond strongly to KIR2DS4 stimulation overriding the lack of licensing.

**Figure 2.**
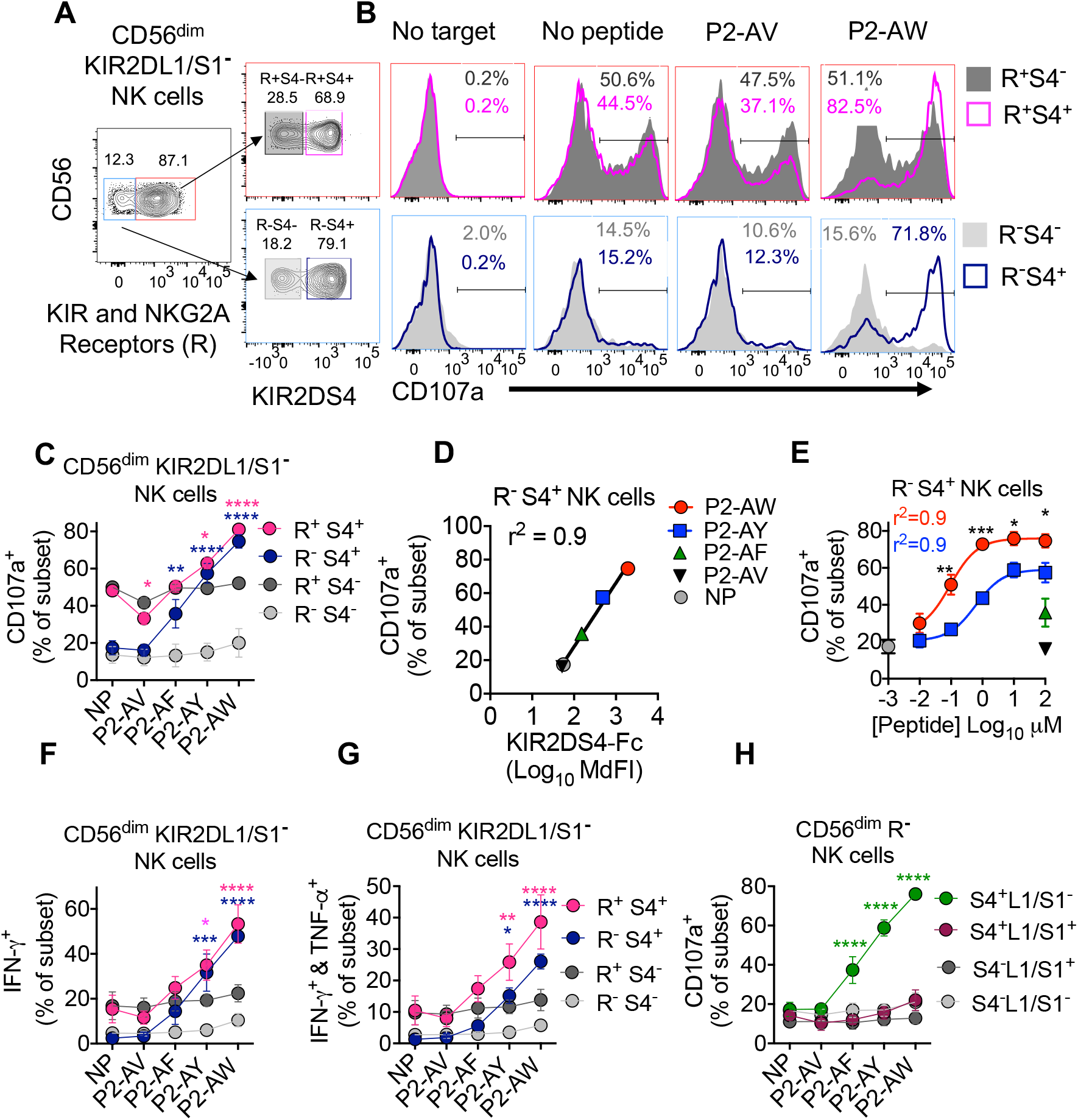
Activation of KIR2DS4^+^ NK cells by peptide:HLA-C*05:01. **(A)** Gating strategy to identify four NK cell subsets by expression of KIR2DS4 (S4) and MHC-I binding receptors (R) within the CD56^dim^ KIR2DL1/S1^−^ NK subset. The R^+^ subset includes NK cells positive for KIR2DL2/L3/S2 (clone GL183), KIR3DL1/3DS1 (clone Z27) and NKG2A (clone Z199). **(B)** Flow cytometry histograms displaying CD107a expression on NK cell subsets defined in **(A)** in response to 221–C*05:01-ICP47 cells loaded with P2-AV, P2-AW, no peptide or no target cells. Data are from one representative experiment out of 3 independent experiments. **(C)** Expression of CD107a on NK cell subsets as in **(B)** in response to 221–C*05:01-ICP47 cells loaded with P2-AV, P2-AF, P2-AY, P2-AW or no peptide. Data are mean and SEM from 3 independent experiments with NK cells from different donors. **(D)** Expression of CD107a on R^−^S4^+^ NK cells from **(C)** is correlated with KIR2DS4-Fc binding to 221–C*05:01-ICP47 cells from Figure. 1D, shown as Log-10 MdFI. Pearson correlation, r^2^ = 0.9, p = 0.0003. **(E)** Expression of CD107a on R^−^ S4^+^ NK cells in response to 221–C*05:01-ICP47 cells loaded with increasing concentrations of P2-AY and P2-AW. R^−^S4^+^ NK cell responses to 221–C*05:01-ICP47 cells loaded with 100 μM of peptides P2-AF and P2-AV or without peptide are also shown. Samples are color coded as in **(D)**. Data are mean and SEM from 3 independent experiments with NK cells from different donors. **(F)** Expression of IFN-γ only and **(G)** co-expression of IFN-γ and TNF-a on NK cell subsets as defined in **(A)** in response to 221–C*05:01-ICP47 cells loaded with P2-AV, P2-AF, P2-AY, P2-AW or no peptide (NP). Data are mean and SEM from 3 independent experiments with NK cells from different donors. (H) Expression of CD107a on R^−^ NK cells in response to 221–C*05:01-ICP47 cells loaded with P2-AV, P2-AF, P2-AY, P2-AW or no peptide. R^−^ NK cells were gated into four populations based on the expression of KIR2DS4 (S4) and KIR2DL1/S1 (L1/S1). Data are mean and SEM from 3 independent experiments with NK cells from different donors. Statistical significance was assessed by two way ANOVA with Dunnett’s multiple comparisons test, *,P < 0,05, **, P < 0.01, ***, P < 0.001, ****, P < 0.0001. In C, G&H, significance is indicated in comparison to NP and is color coded by NK cell subsets. In E, significance is indicated in comparing P2-AW and P2-AY.

There was a strong correlation (r^2^=0.9) between peptide dependent KIR2DS4-Fc binding to HLA-C*05:01 and the functional response of R^−^S4^+^ NK cells in response to the same peptides (Fig. 2C,D). Activation of R^−^S4^+^ NK cells in response to 221-C*05:01-ICP47 cells loaded with P2-AW and P2-AY was dependent on peptide concentration (Fig. 2E). Concentrations of peptide P2-AW lower than 100 nM activated of R^−^S4^+^ NK cells and half-maximal activation was reached with 0.07μM of P2-AW and 1 μM of P2-AY (Fig. 2E). R^−^S4^+^ NK cells in response to 221-C*05:01-ICP47 cells loaded with P2-AW also reached a greater maximum than that was obtained with high concentrations of P2-AY (Fig. 2E). Peptide-specific activation of KIR2DS4^+^ NK cells also induced production of IFN-gamma and TNF-alpha, ascertained by intracellular cytokine staining (Fig. 2F,G and SFig. 2C).

Signaling by ITIM bearing inhibitory receptors dominates over signaling by NK cell activation receptors (Long et al., 2013). The potent activation of NK cells by KIR2DS4, which can override the lack of NK cell licensing, suggested it may be difficult to inhibit. To test whether NK cell activation by KIR2DS4 is regulated by inhibitory receptors, we first re-gated our data from previous degranulation experiments (Fig. 2C). We gated NK cells as CD56^dim^ then gated on R^−^ NK cells without excluding the KIR2DL1/S1 + (EB6) population. These NK cells were then gated into four subsets based on expression of KIR2DS4 (S4) and KIR2DL1/S1 (L1/S1) (Fig. 2H). S4^+^L1/S1^−^ NK cells displayed strong activation in response to 221-C*05:01-ICP47 cells loaded with P2-AF, P2-AY and P2-AW (Fig 2H). In contrast, S4^+^L1/S1+ NK cells displayed no activation even in the response to cells loaded with P2-AW. Because the mAb EB6 binds KIR2DL1 and KIR2DS1, we confirmed these results with two donors who were typed to only carry KIR2DL1 (SFig. 2D,E). As KIR2DS4 is generally expressed by a higher proportion of NK cells than KIR2DL1, many KIR2DS4^+^ NK cells will not be inhibited (SFig. 2F,G). Together, these data show that KIR2DS4^+^ NK cells are potently activated to degranulate and produce cytokines upon engagement of KIR2DS4 and HLA-C*05:01 in a highly peptide selective manner. Activation of KIR2DS4^+^ NK cells was limited to the KIR2DL1^−^ subset due to dominant inhibition by KIR2DL1 + NK cells. R^−^S4^+^ NK cells were unlicensed but potently activated by KIR2DS4 stimulation overriding a lack of NK cell licensing. Therefore, under physiological conditions where the ligands for many NK cell inhibitory receptors are expressed, it is likely that the R^−^S4^+^ NK subset is the major operative subset due to its lack of inhibitory receptor expression.

### Peptide:HLA-C is sufficient to activate KIR2DS4^+^ NK cells

Cross-linking of NK cell activation receptors is not sufficient to trigger resting NK cells, as activation receptors function as synergistic pairs (Bryceson et al., 2005; Bryceson et al., 2006a, b). FcγRIIIA (CD16) is the only exception. To test the requirements for activation of KIR2DS4^+^ NK cells, we performed re-directed antibody mediated degranulation assays with Fc receptor^+^ P815 cells. Minimal NK cell responses were observed when resting NK cells were pre-incubated with mAbs to either NKp46 or 2B4 and then mixed with P815 cells (SFig. 3 A,B). In contrast, strong activation was observed when resting NK cells were pre-incubated with mAbs to both 2B4 and NKp46 (SFig. 3A,B). Remarkably, stimulation of KIR2DS4 alone was sufficient to activate resting NK cell degranulation (SFig. 3 A,B). Thus, activation of resting KIR2DS4^+^ NK cells does not require synergistic stimulation of multiple receptors.

To test whether HLA-C is sufficient to activate KIR2DS4^+^ NK cells, biotinylated HLA-C*05:01 refolded with P2-AV or P2-AW was conjugated to streptavidin Dynabeads. These beads stained strongly with a β_2_M specific mAb (SFig. 3C). KIR2DS4^+^KIR2DL1/S1^−^ (S4^+^L1/S1^−^) NK cells degranulated strongly in response to beads conjugated to HLA-C*05:01 refolded with P2-AW, but not with P2-AV, while KIR2DS4^−^ NK cells were not stimulated by any of the beads (Fig. 3A,B). Activation of KIR2DS4^+^ cells was dependent on bead number (SFig. 3D) and expression of KIR2DL1/S1 on KIR2DS4^+^ NK cells inhibited NK cell activation (SFig. 3E), consistent with our previous results with peptide loaded 221-C*05:01-ICP47 cells (Fig. 2H). Next, HLA-C*05:01 conjugation to Dynabeads was titrated such that the number of HLA-C molecules per bead ranged from 0.5×10^3^ to 10^5^ (Fig. 3C, SFig 3F). The number of HLA-C molecules per bead was estimated using quantitative flow cytometry, by generating a standard curve from beads with known antigen densities (SFig 3F). S4^+^L1/S1^−^ NK cells showed a sharp activation threshold, with half maximal responses at 2000 and 1000 HLA-C molecules per bead for donors 1 and 2, respectively (Fig. 3E). In contrast, S4^+^L1/S1^−^ NK cells were weakly stimulated with beads conjugated with HLA-C*05:01-P2-AV at antigen densities greater than 20,000 HLA-C molecules per bead (Fig. 3D, SFig. 3G). NK cells from a third donor showed a weaker response with a gradual increase in the activation of S4^+^L1/S1^−^ NK cells with increasing HLA-C antigen density and a half maximal response at 8000 HLA-C molecules per bead (Fig. 3E). We concluded that HLA-C alone, immobilized to beads, is sufficient to activate KIR2DS4^+^ NK cells across a range of antigen densities in the absence of ligands for other co-activation receptors or adhesion molecules.

**Figure 3.**
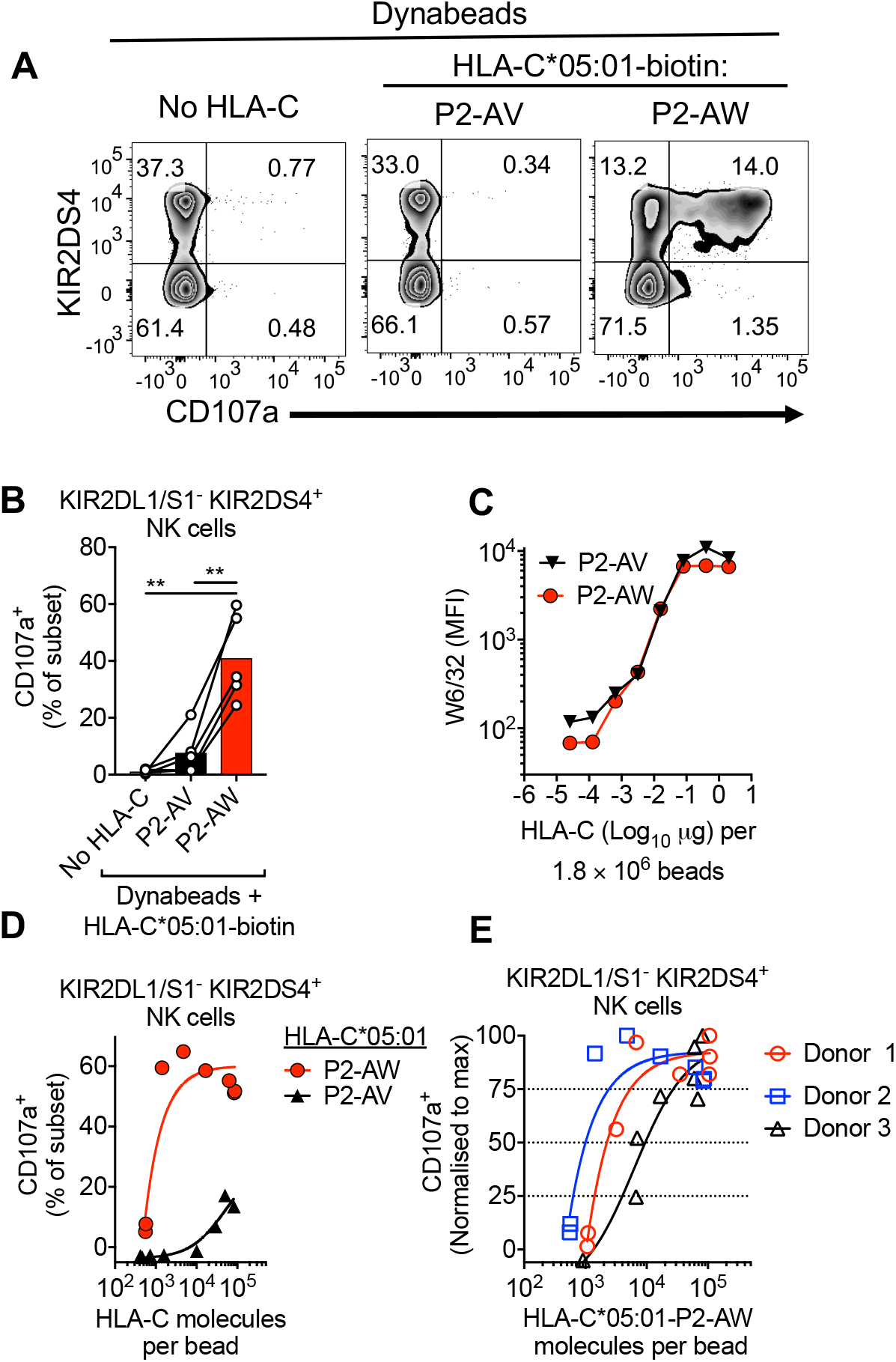
Peptide:HLA-C is sufficient to activate KIR2DS4^+^ NK cells. **(A)** Flow cytometry bi-plots displaying CD107a expression on CD56^dim^ KIR2DL1/S1^−^ NK cells after stimulation with unconjugated Dynabeads and Dynabeads conjugated to biotinylated HLA-C*05:01 refolded with P2-AV or P2-AW. Expression of KIR2DS4 is shown on the y-axis. **(B)** Expression of CD107a on CD56^dim^ KIR2DL1/S1^−^ KIR2DS4^+^ NK cells after stimulation by unconjugated Dynabeads and Dynabeads conjugated to biotinylated HLA-C*05:01 refolded with P2-AV or P2-AW. Data with NK cells from five donors is shown. **(C)** Binding of W6/32 mAb to Dynabeads conjugated to HLA-C*05:01 biotin refolded with P2-AV or P2-AW. Conjugation of HLA-C to 1.8 million Dynabeads was titrated five-fold eight times from 2 μg. One of three independent experiments is shown. **(D)** Expression of CD107a on CD56^dim^ KIR2DL1/S1^−^ KIR2DS4^+^ NK cells after stimulation with Dynabeads bearing increasing densities of biotinylated HLA-C*05:01 refolded with P2-AV or P2-AW. Antigen density was calculated quantitative flow cytometry. Data from donor 2 is shown. **(E)** Expression of CD107a on CD56^dim^ KIR2DL1/S1^−^ KIR2DS4^+^ NK cells after stimulation with Dynabeads bearing increasing antigen densities of biotinylated HLA-C*05:01 refolded with P2-AW. Data from three donors were normalized to the maximum value of CD107a+ for each donor. Maximum responses were for Donor 1 = 42%, Donor 2 = 65%, Donor 3 = 32%. Statistical significance was assessed by one way ANOVA with Tukey’s multiple comparisons test, **, P < 0.01.

### Functional presentation of endogenous P2-AW peptide by HLA-C*05:01 to KIR2DS4^+^ NK cells

Our experiments thus far demonstrated that KIR2DS4^+^ NK cells can recognize peptides presented by HLA-C*05:01 in a highly selective manner. For these experiments, the peptides presented by HLA-C*05:01 have been dominated by single peptide sequences, in the form of recombinant HLA-C refolded with peptide or TAP-deficient cells loaded with peptide. While useful experimental tools, under physiological conditions HLA-C*05:01 does not present single peptides, but many peptides of different sequences (Di Marco et al., 2017; Kaur et al., 2017; Sim et al., 2017). To test whether KIR2DS4^+^ NK cells could recognize peptide in TAP sufficient cells where HLA-C*05:01 presents many different peptides, we expressed the peptides P2-AW or P2-AV in 221-C*05:01 cells using a retrovirus (Fig. 4A, SFig. 5A) (Gejman et al., 2018). Transduced cells were marked by expression of mCherry, and after drug selection both mini-genes were expressed in over 90% of cells (SFig. 5B). In degranulation assays, R^−^S4^+^ NK cells responded weakly to both 221-C*05:01 cells and those transduced with P2-AV (Fig. 4B). In contrast, R^−^S4^+^ NK cells exhibited enhanced degranulation in response to 221-C*05:01 cells expressing the P2-AW peptide (Fig. 4B). This activation was KIR2DS4 specific, as R^−^S4^−^ NK cells exhibited low responses to both 221-C*05:01 cells and those transduced with P2-AV or P2-AW. Thus, KIR2DS4 can detect a stimulatory peptide, presented by HLA-C*05:01 in TAP sufficient cells that present many different peptide sequences.

**Figure 4.**
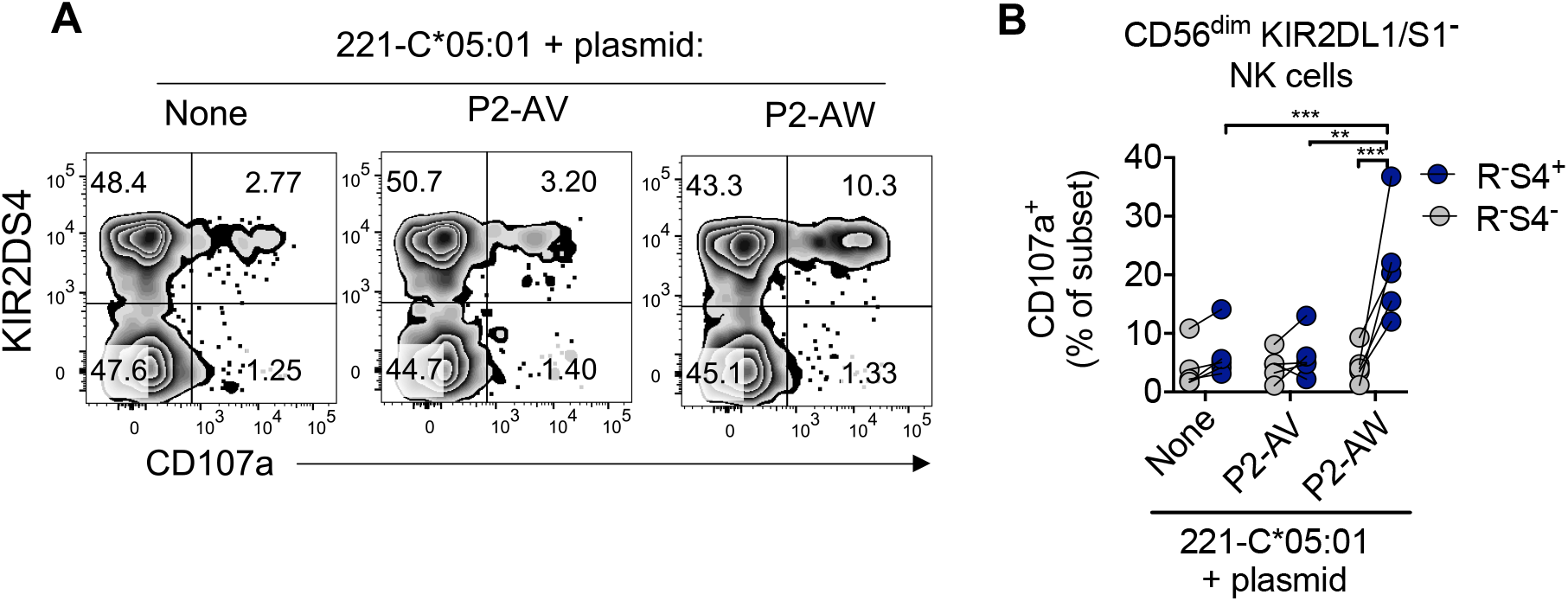
Functional presentation of endogenous P2-AW peptide by HLA-C*05:01 to KIR2DS4^+^ NK cells. **(A)** Expression of CD107a on KIR2DS4^+^ and KIR2DS4^−^ NK cell subsets in response to 221–C*05:01 cells transduced with plasmids encoding P2-AV or P2-AW. NK cells were gated as CD56^dim^ KIR2DL1/S1^−^ R^−^ as in Fig. 2A. One representative experiment of five is shown. **(B)** Expression of CD107a on NK cell subsets as in **(A)** in response to 221–C*05:01 cells transduced with plasmids encoding P2-AV or P2-AW. Data from five independent experiments are shown. Statistical significance was assessed by two way ANOVA with Sidak’s multiple comparisons test, **, P < 0.01, ***, P < 0.001.

### A ‘self’ peptide eluted from HLA-C*05:01 carrying Tryptophan at position 8 is a KIR2DS4 epitope

We reasoned that peptides containing Trp at p8 may be enriched for KIR2DS4 binding peptides, as in the context of P2-AY, Trp at p8 conferred stronger binding of KIR2DS4 to HLA-C*05:01 than Tyr or Phe (Fig. 1). Peptides eluted from HLA-C*05:01 contain a low frequency of Trp (SFig. 5), which was similar to peptides eluted from other HLA-C allotypes (Di Marco et al., 2017; Kaur et al., 2017). Tryptophan is rare at p8 for all HLA-C allotypes, ranging from 0-2.6%, and is found at a frequency of 0.6% in HLA-C*05:01 peptides (SFig. 5B). The scarcity of peptides containing Trp at p8 presented by HLA-C*05:01 may explain the lack of KIR2DS4-Fc binding to 221-C*05:01 cells.

We synthesized and tested the 12 peptides with Trp at p8 that had been eluted from HLA-C*05:01 for KIR2DS4 binding and activation of KIR2DS4^+^ NK cells. One strong and one weak KIR2DS4 binding ‘self’ peptides were identified (Fig. 5). The KIR2DS4 weak binding peptide was TM9SF4_323-311_ (MSDVQIHWF). TM9SF4 is a member of the transmembrane superfamily 9, a highly conserved family across evolution with roles in cell adhesion, phagocytosis and autophagy (Paolillo et al., 2015). The KIR2DS4 strong binding ‘self’ peptide was HECTD1_1131-1139_ (SNDDKNAWF). The Homologous to the E6-AP Carboxyl Terminus (HECT) domain E3 ubiquitin ligase 1 (HECTD1) has been linked to cholesterol export from macrophages (Aleidi et al., 2018) and suppression of epithelial-to-mesenchymal transition in cancer metastasis (Duhamel et al., 2018). Both TM9SF4 and HECTD1 are expressed in all tissues at the RNA and protein levels (Uhlen et al., 2015). That only two of 12 peptides with Trp at p8 bound KIR2DS4 emphasized the high peptide specificity of KIR2DS4 binding to HLA-C*05:01 and that Trp is not sufficient to identify KIR2DS4 binding peptides.

**Figure 5.**
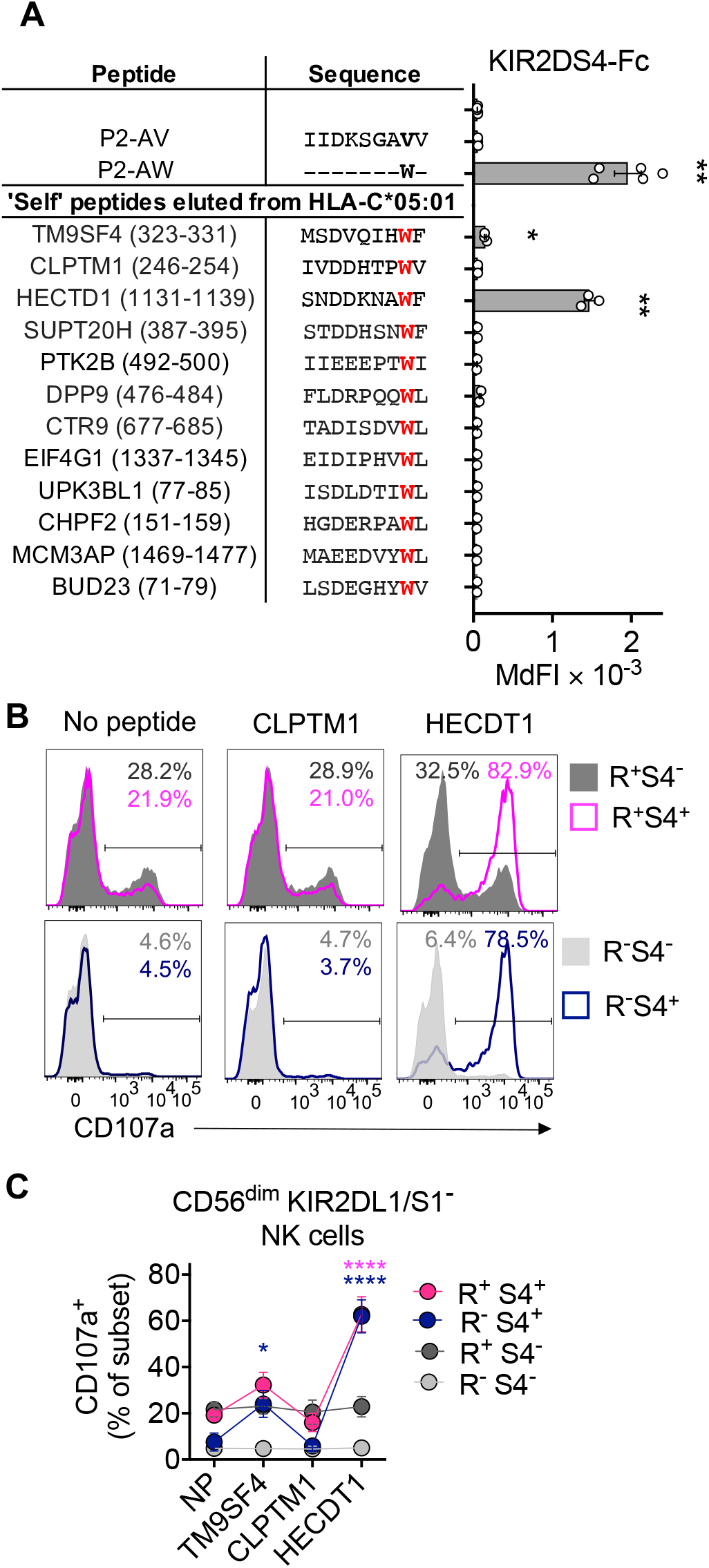
A ‘self’ peptide eluted from HLA-C*05:01 with tryptophan at position 8 is a KIR2DS4 epitope. **(A)** KIRD2S4-Fc binding to 221-C*05:01-ICP47 cells loaded with each one of 12 endogenous ‘self’ peptides carrying Trp at position 8, compared to P2-AV, P2-AW or no peptide. The 12 ‘self’ peptides were identified from two studies, which eluted and sequenced a total of 1674 unique 9mer peptides from purified HLA-C*05:01. TM9SF4_323-11_ and HECTD1_1131-1139_ were identified in both studies. The amino acid sequence of each ‘self’ peptide and position within the protein of origin is listed. **(B)** Flow cytometry histograms displaying CD107a expression on CD56^dim^ KIR2DL1/S1^−^ NK cell subsets in response to 221-C*05:01-ICP47 cells with loaded with ‘self’ peptides from proteins CLPTM1 and HECDT1 or no peptide. CD56^dim^ KIR2DL1/S1^−^ NK cells were gated as in Fig 2A. **(C)** Expression of CD107a on NK cell subsets as in **(B)**. Mean and SEM from 3 independent experiments with NK cells from different donors are shown. Statistical significance was assessed by one way ANOVA with Tukey’s multiple comparisons test **(A)** and two way ANOVA with Dunnett’s multiple comparisons test **(C)**, *,P < 0,05, **, P < 0.01, ****, P < 0.0001. Significance is indicated in comparison to NP and is color coded by NK cell subsets.

### A conserved KIR2DS4 epitope derived from Recombinase A is shared by hundreds of species of bacteria

To explore whether KIR2DS4 recognizes pathogen-derived peptides, we searched the proteomes of prokaryotes for sequences similar to P2-AW. P2-AW (IIDKSGAWV) showed remarkable homology to residues 283-291 of Recombinase A (RecA) of *Helicobacter fennelliae* (IIDKSGAWI; Fig. 6A). RecA is the prototypical DNA recombinase and is essential for DNA damage repair by homologous recombination (Lusetti and Cox, 2002). A previous study aligned the sequences of RecA proteins from 63 species of bacteria and found amino acid sequence similarity ranged from 43% to 100% (Karlin and Brocchieri, 1996). Residues 283-291 are highly conserved, and in particular, G_288_ is conserved in all 63 species, W_290_ is found in 61 out of 63 species and acidic residues at position 285 are found in 40 out of 63 species (SFig. 6A). A logo-motif for RecA_283-291_ from these 63 species shown in Figure 6B.

**Figure 6.**
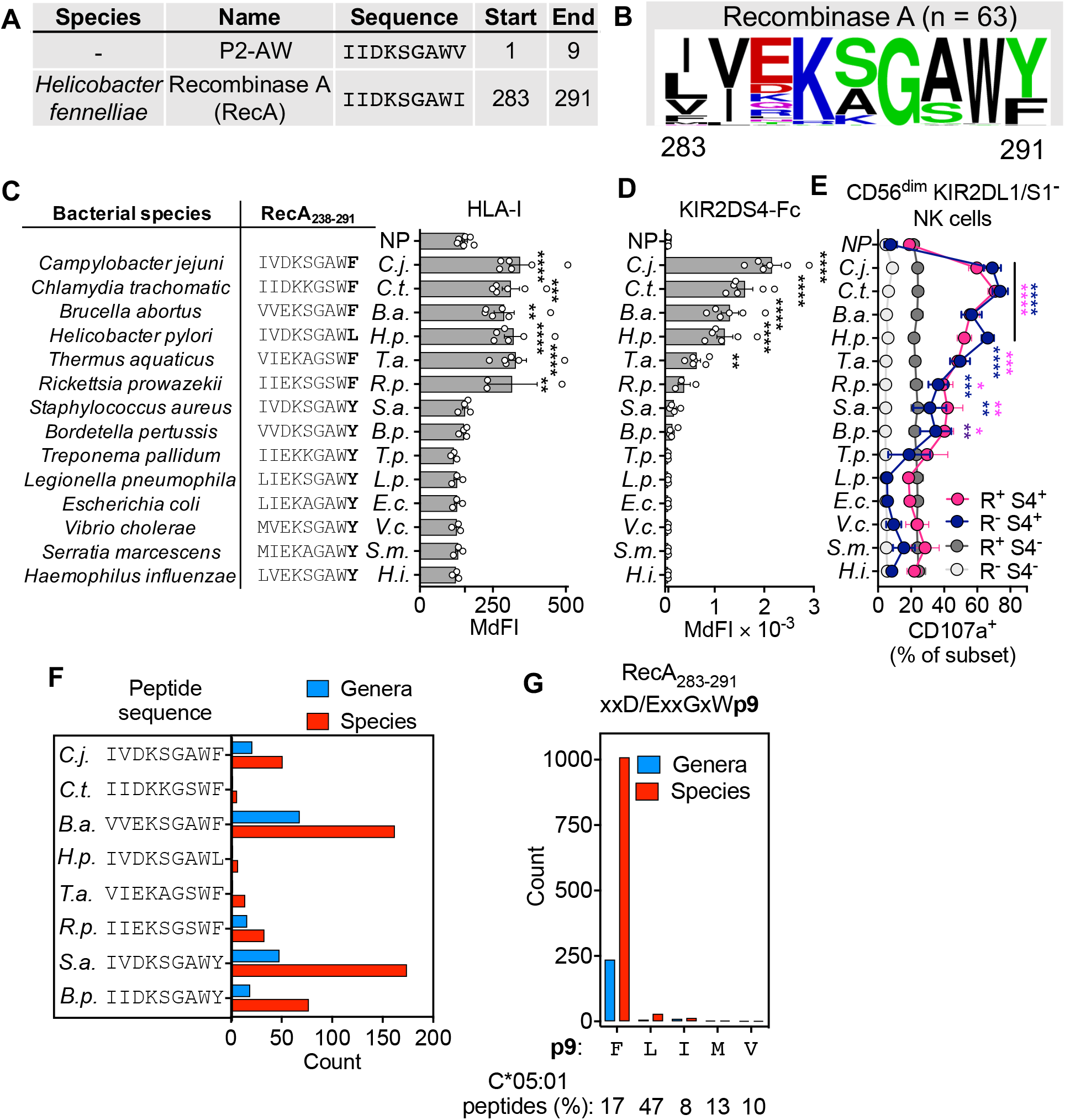
A KIR2DS4 epitope in a conserved recombinase A sequence shared by hundreds of bacterial species. **(A)** Amino acid sequence alignment of peptide P2-AW and Recombinase A (RecA)_283-291_ sequence from *Helicobacter fennelliae*. **(B)** Peptide sequence motif present in RecA_283-291_ sequences from 63 species of bacteria. **(C)** Stabilization of HLA-I expression on 221-C*05:01-ICP47 cells incubated overnight at 26°C with 100 μM of 14 different RecA_283-291_ peptides. Recombinase A_283-291_ peptide sequences are from 14 species of bacteria. Data shown as median fluorescence intensity (MdFI), n=3-5. **(D)** KIR2DS4-Fc binding to 221-C*05:01-ICP47 cells incubated with peptides as shown in **(C)**. Data shown as MdFI, n=3-5. **(E)** Expression of CD107a on CD56^dim^ NK cell subsets in response to 221-C*05:01-ICP47 cells loaded with peptides as in **(C)**. CD56^dim^ KIR2DL1/S1^−^ NK cells were gated as in Fig. 2A. Data are mean and SEM from 3 independent experiments with NK cells from different donors. **(F)** Number of genera and species of bacteria with RecA_283-291_ peptide sequences identical to those that activate KIR2DS4^+^ NK cells as in F. **(G)** Number of genera and species of bacteria with Recombinase A_283-291_ peptide sequences with the motif xxD/ExxGxWp9, x = any residue, p9 = C-terminal residue. The frequency of F, L, I, M and V at the c-terminus of 9mer peptides eluted from HLA-C*05:01 is also shown. Statistical significance was assessed by one way (D&E) and two way **(F)** ANOVA, *,P < 0,05, **, P < 0.01, ***, P < 0.001, ****, P < 0.0001. Significance is indicated in comparison to NP and in F is color coded by NK cell subsets.

We tested 14 RecA_283-291_ peptides with different sequences, each containing Asp or Glu at position 3, a critical anchor residue for HLA-C*05:01. Due to the high level of conservation, these 14 peptides cover more than 14 species of bacteria; for example, RecA_283-291_ from *Yersinia pestis* is identical to that of *Escherichia coli*. RecA_283-291_ peptides with Phe or Leu at the C-terminus stabilized HLA-C*05:01 well, bound KIR2DS4 and activated KIR2DS4^+^ NK cells (Fig. 6C-F). Epitopes derived from the pathogens *Chlamydia trachomatic, Campylobacter jejuni, Brucella abortus* and *Helicobacter pylori* bound KIR2DS4 the strongest and conferred potent stimulation of KIR2DS4^+^ NK cells (Fig. 6C-F). The epitopes from *Thermus aquaticus* and *Rickettsia prowazekii* conferred an intermediate level of KIR2DS4 binding, while epitopes from *Stapylococcus aureus* and *Bordetella pertusis* conferred weak KIR2DS4 binding (Fig. 6C-F). Of the 14 peptides, those with Tyr at the C-terminus were poor ligands for HLA-C*05:01 and conferred low HLA-I stabilization and little or no KIR2DS4 binding (Fig. 6C-F). This is consistent with the very low frequency of peptides eluted from HLA-C*05:01 that use Tyr as a C-terminal anchor (SFig. 6B).

To evaluate how many species of bacteria may contain KIR2DS4 epitopes, we downloaded all bacterial RecA sequences from the protein database (https://www.ncbi.nlm.nih.gov/protein). We identified over a hundred bacterial species with RecA_283-291_ sequences identical to those that bound KIR2DS4 (Fig. 6G). Of the four sequences that bound KIR2DS4 the strongest, those identical to the *Brucella abortus* sequence were the most frequent. This included the sequence from *Brucella melitensis*, another pathogenic species of the *Brucella* genus (de Figueiredo et al., 2015). Taking a broader approach, we generated a RecA_283-291_ motif (xxD/ExxGxW**p9**) accounting for the essential role of acidic resides at position 3 for binding HLA-C*05:01, the high conservation of p6 G and the importance of p8 Trp for binding KIR2DS4 (Fig. 6G). We allowed for any amino acid at all other positions (x) expect p9. To focus our analysis on peptides predicted to be presented well by HLA-C*05:01, we counted only RecA_283-291_ sequences where Phe, Leu, Ile, Met or Val at p9, all common C-terminal anchors for HLA-C*05:01 peptides (SFig. 6B). Using this motif, we identified over one thousand different bacterial species that contain RecA_283-291_ sequences that have the potential to be presented by HLA-C*05:01 and bind KIR2DS4 (Fig. 6G). The majority of species containing RecA_283-291_ sequences predicted to bind KIR2DS4 were from Proteobacteria and within the Proteobacteria the most common Order was Rhizobiales (SFig. 6C,D). All bacterial species containing RecA sequences predicted to be presented by HLA-C*05:01 and recognized by KIR2DS4 are shown in Supplementary Table 1. Together our data suggest the possibility that HLA-C*05:01^+^ individuals expressing KIR2DS4 may have an evolutionary advantage in bacterial immunity, through the ability of their NK cells to recognize RecA epitopes presented by HLA-C*05:01.

### Allele frequency of full-length KIR2DS4 is inversely correlated with the frequency of HLA-C*05:01

The allele frequency of functional KIR2DS4-fl (*KIR2DS4*001*) is positively correlated with HLA-A*11, a previously identified KIR2DS4 ligand (Graef et al., 2009). We therefore examined the allele frequency of HLA-C*05:01 in 10 populations where the percentage of individuals carrying KIR2DS4-fl (*KIR2DS4*001*) and KIR2DS4-del (*KIR2DS4*003*) is known (http://www.allelefrequencies.net/) (Robinson et al., 2015). There was a strong negative correlation (r= −0.83, r^2^=0.7, p=0.003) with the allele frequency of *HLA-C*05:01* and the percentage of individuals carrying *KIR2DS4*001* (Fig. 7A). Conversely, there was a positive correlation (r=0.64, r^2^=0.4, p=0.04) with the allele frequency *HLA-C*05:01* and the percentage of individuals carrying *KIR2DS4*003* (Fig. 7B). The highest frequency of *HLA-C*05:01* is found in those with European Caucasian ancestry represented by two populations; one from Northern Ireland (0.13) and the other from the USA (0.09; Fig. 7A). The percentage of individuals carrying *KIR2DS4*001* in these two populations was lower than 45% (Fig. 7A). In contrast, populations with very low frequency of *HLA-C*05:01*, shown here by two Chinese populations (0.0018) and a South African population (0.008), had greater than 75% of individuals carrying *KIR2DS4*001* (Fig. 7A). We saw similar results using data from 9 populations with high resolution KIR allele typing where the allele frequencies of all three genotypes KIR2DS4-fl, KIR2DS4-del and KIR2DS4 negative were known (SFig. 7A,B). The allele frequency of *HLA-C*05:01* was inversely correlated with the frequency of KIR2DS4-fl alleles (r= −0.85, r^2^=0.7,p=0.004), positively correlated with the frequency of KIR2DS4-del alleles (r=0.95, r^2^=0.9, p=0.0004) and the frequency of the KIR2DS4 negative genotype combined with the frequency of KIR2DS4-del alleles (r=0.85, r^2^=0.7, p=0.004; SFig. 7A,B). This effect appeared to be unique to HLA-C*05:01 as the allele frequencies of other KIR2DS4 ligands HLA-C*04:01 and HLA-C*16:01 (Graef et al., 2009), showed no correlation with the percentage of individuals carrying *KIR2DS4*001* or *KIR2DS4*003* (Fig. 7C-F). Therefore, populations with a higher frequency of *HLA-C*05:01* have lower frequencies of the functional receptor KIR2DS4-fl.

**Figure. 7.**
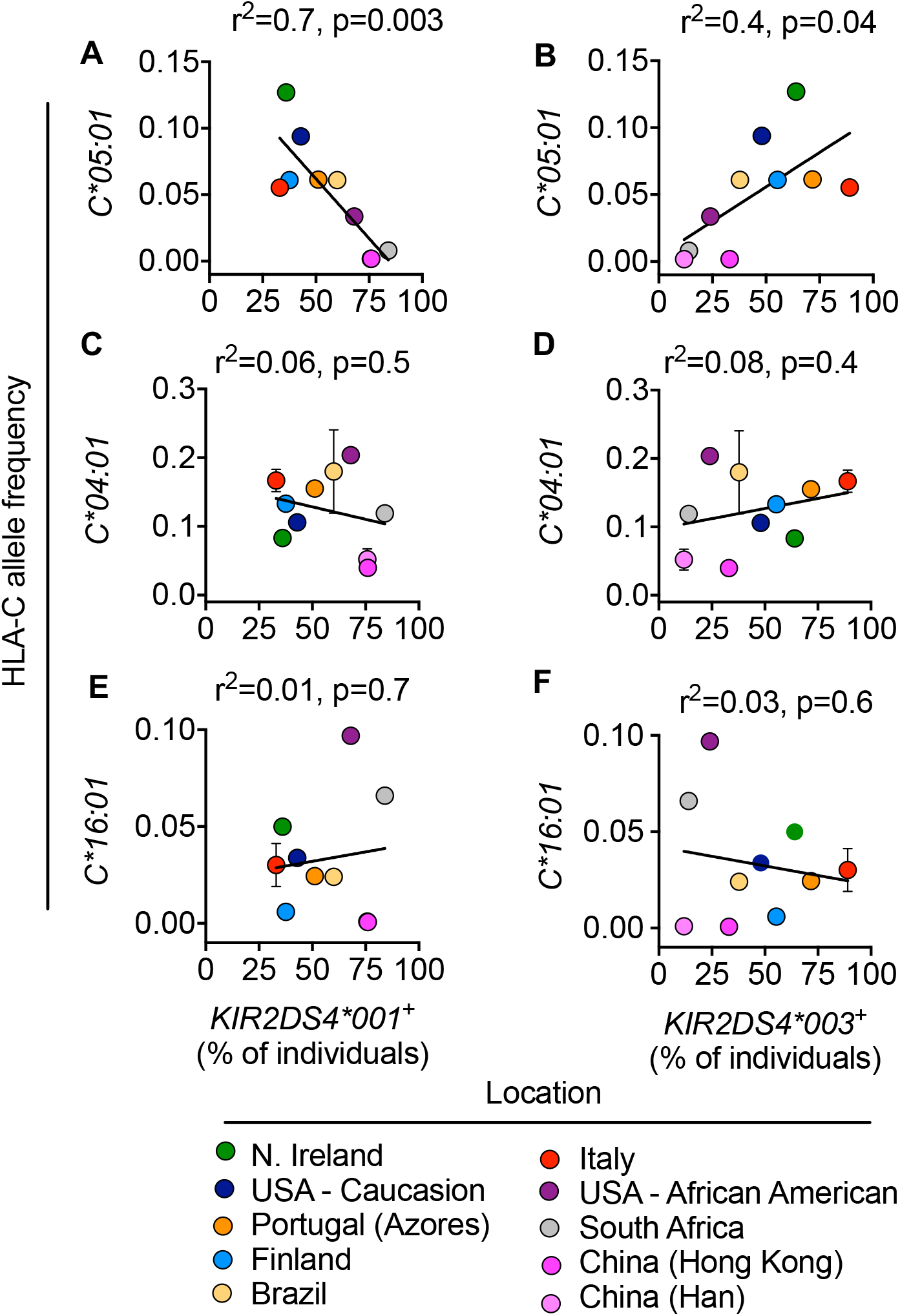
Functional KIR2DS4 allele is inversely correlated with HLA-C*05:01. Allele frequency of *HLA-C*05:01* **(A,B)**, *HLA-C*04:01* **(C,D)** and *HLA-C*16:01* **(E,F)** correlated with the percentage of individuals who are *KIR2DS4*001*^+^ **(A,C,E)** and *KIR2DS4*003*^+^ **(B,D,F)**, in 10 populations. Data are from www.allelefrequencies.net.

## Discussion

The concept of an innate receptor with specificity for one form of a highly variable antigen, such as an MHC-peptide complex, is counterintuitive. And yet, members of the killer-cell Ig-like receptor (KIR) family bind to HLA class I allotypes with various degrees of selectivity for peptides (Malnati et al., 1995; Rajagopalan and Long, 1997; Sim et al., 2017). In the case of the activating members of the KIR family, for which ligands are less well defined, a reasonable hypothesis was that they serve to recognize conserved features of modified ‘self’. Here we describe an extreme case of peptide selectivity of an activating KIR, which is restricted by HLA-C*05:01. Peptides that promote binding of KIR2DS4 carried a Trp at position 8 of 9-mer peptides. Recognition of this peptide–HLA-C*05:01 complex by KIR2DS4^+^ NK cells induced potent activation of NK cells, and could even activate unlicensed NK cells. Interestingly, optimal peptides for activation of KIR2DS4^+^ NK cells identified in this study included an epitope that is highly conserved in the essential protein RecA expressed by hundreds of bacterial species, suggesting KIR2DS4 may have evolved to contribute to bacterial innate immunity.

RecA is essential for repair of damaged DNA by homologous recombination and is critical for the DNA damage response in *E. coli* (Lusetti and Cox, 2002). The epitope in the bacterial protein RecA for KIR2DS4 binding contains Trp at p8 and this position (residue 290 in *E. coli*) is conserved across bacteria, including many human pathogens. Tryptophan is a rare amino acid in general (1.3% of the human proteome), which is reflected in its low representation in peptides eluted from HLA-C (Di Marco et al., 2017; Kaur et al., 2017). Only 0.6% of peptides (n = 2035) eluted from HLA-C*05:01 contained Trp at p8. Out of 12 such peptides, only one bound KIR2DS4, providing an explanation for the lack of KIR2DS4 binding to, or activation of KIR2DS4^+^ NK cells by 221–C*05:01 cells. The low abundance of peptides including Trp at p8 provides the opportunity to detect similar but more abundant peptides, perhaps modified peptides or peptides derived from pathogens. We propose that KIR2DS4 is not specific for non-self but for ‘rare self’, which provides the opportunity to detect abundant foreign peptides carrying a ‘rare self’ epitope.

The major function of inhibitory KIR is recognition of ‘self’ to inhibit NK cell activation and dictate NK cell licensing (Long et al., 2013). Inhibitory KIR exhibit limited peptide selectivity and bind HLA class I in the context of many biochemically diverse peptides. However, differences in peptide selectivity exist between KIRs. Recently, we compared KIR2DL1 binding to HLA-C*05:01 and KIR2DL2/3 binding to HLA-C*08:02 in the presence of the same peptides. HLA-C*05:01 and HLA-C*08:02 differ only by the two amino acids that define the C1 and C2 epitopes and could be loaded with the same peptides. While KIR2DL1 bound HLA-C*05:01 in the presence of 24 out of 28 peptides, KIR2DL2 and KIR2DL3 bound HLA-C*08:02 in the presence of only 13 and 11 of the same peptides, respectively (Sim et al., 2017). Position 44 of KIR2DL1 (Met) and KIR2DL2/3 (Lys) dominates in determining KIR specificity for C2 and C1 allotypes, respectively (Boyington et al., 2000; Fan et al., 2001). However, Lys44 receptors exhibit cross-reactive binding to C2 allotypes (Moesta et al., 2008; Winter et al., 1998). KIR2DS4, which carries Lys44, has the capacity to bind a small subset of C1 and C2 allotypes. The high peptide selectivity of KIR2DS4 is reminiscent of cross-reactive KIR binding, where KIR2DL2/3 bound HLA-C*05:01 and KIR2DL1 bound HLA-C*08:02 in the presence of 5 and 2 out of 28 of the same peptides, respectively (Sim et al., 2017). KIR2DS4 was even more peptide selective than this, binding none of the same 28 peptides previously studied and only in the presence of the few peptides described here.

Examples of pathogenic bacteria that carry an epitope within RecA for HLA-C*05:01-restricted binding of KIR2DS4 include: *Helicobacter pylori*, an infectious agent associated with peptic ulcers and gastric cancers (Suerbaum and Michetti, 2002); bacteria of the *Brucella* genus, which cause a zoonotic infection called brucellosis (Skendros et al., 2011); *Campylobacter jejuni*, a major cause of gastroenteritis with an incidence as common as that caused by *Salmonella* infections (Marder et al., 2017); and *Chlamydia trachomatis*, which is the most common sexually transmitted infection with over 90 million cases annually worldwide (Brunham and Rey-Ladino, 2005). There are bacterial species with a RecA sequence that lacks an Asp or Glu at position 3, which is required for peptide binding to HLA-C*05:01 (Di Marco et al., 2017; Kaur et al., 2017; Sim et al., 2017). Such species include *Mycobacterium tuberculosis* which carries the conserved Trp at p8 but has an Arg at position 3 (LI<u>R</u>KSGAWF). Additionally, other species such as *E. coli* and *Yersinia pestis* carry a Tyr at p9, which is a poor C-terminal anchor for HLA-C*05:01. It is possible that other HLA-C allotypes may also present these RecA peptides by accommodating other amino acids at p3 or p9, expanding the coverage of pathogenic bacteria detected by KIR2DS4^+^ NK cells.

KIR2DS4 is a strong activator of NK cells and was sufficient on its own, to elicit functional responses by KIR2DS4^+^ NK cells. Using an assay with recombinant HLA-C*05:01 carrying an optimal Trp8 peptide conjugated to beads, we determined that the number of HLA-C molecules required for half maximal activation of KIR2DS4^+^ NK cells to be from 1000-8000 suggesting the number of molecules required to activate KIR2DS4^+^ NK cells may be within a physiological range. Furthermore, the presence of ligands on target cells for adhesion molecules and co-activation receptors on NK cells may reduce the number of HLA-C:peptide complexes on human cells required to activate KIR2DS4^+^ NK cells. As a tool to determine whether an endogenous peptide carrying an epitope for HLA-C*05:01-restricted binding of KIR2DS4 could be presented at the plasma membrane and detected by KIR2DS4^+^ NK cells, we expressed peptide P2-AW with a retrovirus. KIR2DS4^+^ NK cells were activated by 221–C*05:01 cells only after endogenous expression of the right peptide. Thus, KIR2DS4^+^ NK cells can detect antigen presented by TAP sufficient cells, presumably in the context of many other peptides that do not bind KIR2DS4. The strong activation signals transmitted by KIR2DS4 overcome the lack of licensing but were still controlled through co-engagement of inhibitory KIR2DL1 on NK cells. NK cell activation by KIR2DS4 must occur in scenarios where HLA-C*05:01 is expressed, where MHC-I is not downregulated unlike ‘missing-self’ scenarios. It is highly likely that KIR2DS4^+^ NK cell subsets that co-express inhibitory receptors will be inhibited, depending on the donor KIR and HLA-I genotypes. However, we demonstrated that even those KIR2DS4^+^ NK cells that lack inhibitory receptors (R^−^S4^+^) can be potently activated via KIR2DS4. Under such scenarios, we propose that this subset (R^−^S4^+^) would be most potently activated.

As NK cells are potently activated by KIR2DS4, overstimulation could have negative consequences, such as inflammation and autoimmunity. The negative correlation between the frequency of HLA-C*05:01 and KIR2DS4-fl alleles in different geographically defined populations, suggests there are detrimental consequences for populations with high frequencies of this receptor and ligand pair, indicative of balancing selection. It demonstrates a genetic interaction between them and suggests that the KIR2DS4-del alleles may have originated in populations with high frequency of HLA-C*05:01. The highest frequency of the HLA-C*05:01 allele is in European populations, including Northern Ireland, where the KIR2DS4-del allele was first described (Maxwell et al., 2002; Middleton et al., 2007; Robinson et al., 2015). This is the first evidence for balancing selection between an activating KIR and its ligand, which is reminiscent of examples of balancing selection in populations with high frequencies of inhibitory KIR ligands (Parham et al., 2012). For example, the frequency of C2 allotypes is very high in the KhoeSan tribes of southern Africa, where novel KIR2DL1 alleles were discovered that have lost C2 specificity or lost the capacity to signal (Hilton et al., 2015b). Similarly, in the Yucpa tribe of South America, where C1 frequency is highest worldwide, novel KIR2DL3 alleles were discovered that encode receptors with weaker avidity for C1 allotypes (Gendzekhadze et al., 2009).

Activating KIR receptors are rapidly evolving and differ even among higher primate species (Abi-Rached and Parham, 2005). Activating KIR evolved from their inhibitory counterparts and co-opted existing evolutionarily conserved signaling adaptor molecules like DAP12 (Abi-Rached and Parham, 2005). However, there is no ancestral lineage of activating KIR genes and they appear to undergo periods of positive selection followed by negative selection. As such, many activating KIR, including *KIR2DS4* are not fixed on either KIR A or B haplotypes suggesting they are under current selection. Beneficial effects of a novel activating KIR could include resistance to pathogens and improved reproductive success, while negative effects could include autoimmunity and too high a birth weight (Abi-Rached and Parham, 2005). Human pregnancy is a physiological process impacted by KIR genes, including KIR2DS4 (Kennedy et al., 2016; Parham and Moffett, 2013). Human birth weight is under balancing selection as babies born too big or too small are less likely to survive, and combinations of KIR and their ligands associate with both ends of this spectrum (Hiby et al., 2014; Parham and Moffett, 2013). Given that KIR2DS4 protects from pre-eclampsia, a disorder or insufficient blood supply to the fetus, it is possible that a high frequency of KIR2DS4 and HLA-C*05:01 had a negative effect in those populations because of too high a birth weight. This kind of balancing effect has been observed for KIR2DS1 and HLA-C2, where this combination of C2 and activating receptor is associated with increased birth weight, while the combination of C2 and inhibitory receptor is associated with low birth weight and increased risk of pre-eclampsia (Hiby et al., 2014; Parham and Moffett, 2013).

An NK cell activating receptor with high selectivity for pathogen-derived peptides is not without precedent. The CD94:NKG2C receptor was recently shown to recognize peptides derived from the UL40 protein of human cytomegalovirus (HCMV), presented by HLA-E (Hammer et al., 2018). Recognition of these peptides is thought to drive the formation of adaptive NK cells in HCMV infected individuals (Hammer et al., 2018). Furthermore, a recent study discovered that KIR2DS2, another activating KIR, recognizes HLA-C-bound peptides derived from a conserved region of the NS3 helicase of the Flavivirus family (Naiyer et al., 2017). This family includes several pathogens such as Hepatitis C, Ebola, Zika and West Nile viruses. It is likely that KIR2DS2^+^ and CD94:NKG2C^+^ NK cells participate in immune defense through direct recognition of virus-infected cells. Here, we demonstrated that KIR2DS4 is a *bona fide* HLA-C binding receptor that may have evolved to play a protective role in immune defense against bacteria. In the context of bacterial infections, the stimulation of IFN-γ production by KIR2DS4^+^ NK cells is likely to contribute to clearance of bacterial pathogens. The first line of defense against invading bacteria are phagocytic, antigen-presenting cells (APC) such as monocytes, macrophages and dendritic cells. NK cells co-operate with these cell types to produce IFN-γ via cell contact-dependent mechanisms and IL-12 production (Ferlazzo et al., 2003; Gerosa et al., 2002; Walzer et al., 2005). It is possible that KIR2DS4^+^ NK cells could directly recognize bacterially infected cells, indeed *C. trachomatis, C. jejuni* and *B. abortus* are intracellular bacteria (Brunham and Rey-Ladino, 2005; Skendros et al., 2011; Watson and Galan, 2008). An additional scenario is that KIR2DS4^+^ NK cells could be activated upon interaction with APCs that present RecA epitopes on HLA-C to produce IFN-γ early during infection, and facilitate a Th1 response necessary to clear the bacterial infection. In this case, the RecA epitope would be presented by HLA-C through crosspresentation, the process whereby exogenous antigens enter the MHC-I pathway (Blander, 2018). The DC subset associated with efficient cross-presentation consists of classical DC1 (cDC1) (Joffre et al., 2012). NK cells interact with DCs, and the interplay between cDC1 and NK cells was shown recently to be important for anti-tumor responses and associated with greater responses to checkpoint blockade therapy (Barry et al., 2018; Bottcher et al., 2018). Notably, NK cells produced CCL5 and XCL1 which facilitated cDC1 recruitment to the tumor site (Bottcher et al., 2018). Thus, in addition to IFN-γ, NK cells may also contribute to enhanced bacterial clearance through promotion of cDC and T cell interactions. Disease association studies of KIR2DS4 and HLA-C*05:01 with outcome of bacterial infections will be needed to test this idea further.

## Methods

Cell lines. 721.221 (221) cells and 221 cells expressing HLA–C*04:01 were used (provided by J. Gumperz and P. Parham, Stanford University, USA). The 221 cells expressing HLA-C*05:01 (221–C*05:01) and those expressing the TAP inhibitor ICP47 (221–C*05:01–ICP47) were previously described (Sim et al., 2017). All 221 cells were cultured in IMDM (Gibco) supplemented with 10% FCS.

### Peptide HLA-C stabilization assays

Peptide stabilization of HLA-C was assessed by flow cytometry largely as described (Sim et al., 2017). 10^5^ cells were incubated overnight at 26°C with 100 μM of synthetic peptide. The following day cells stained with APC HLA-I mAb (W6/32, Biolegend) at 4°C. Peptides were synthesized by Genscript (USA).

### KIR-Fc binding assay

KIR-Fc binding to cell lines and peptide-loaded cells was assessed by flow cytometry largely as described (Sim et al., 2017). KIR2DL1*001-Fc, and KIR2DS4*001-Fc (R&D, 1844-KR, 1847-KR) were conjugated to protein-A Alexa Flour 647 (Invitrogen) by overnight incubation at 9:1 (molar) at 4°C, then diluted to 3.6 μg/ml (KIR2DL1*001-Fc) and 9 μg/ml (KIR2DS4*001-Fc) in PBS + 2% fetal calf serum (FCS). 10^5^ cells were placed in 96 flat well plates, resuspended in 25 μl of KIR-Fc and incubated at 4°C for 1 hr. For peptide loaded cells, KIR-Fc binding was assessed after overnight incubation of cells at 26°C with 100 μM of synthetic peptide. KIR-Fc binding to 221 cells or protein A-Alexa Flour 647 alone were used to baseline values for KIR-Fc binding. Cells were washed with PBS three times and data acquired by flow cytometry.

### NK cell functional assays

NK cell degranulation assays were carried out largely as described (Sim et al., 2017). Primary, resting NK cells were isolated by negative selection from PBMCs and were greater than 95% CD56+, less than 5% CD3+ (EasySep NK cell Isolation Kit, Stem Cell, USA). Donors were screened for the presence of KIR2DS4 with the mAb JJC11.6 (Miltenyi Biotec). 10^5^ resting NK cells were mixed at 1:1 with 221 cell lines or peptide loaded target cells in presence of 1 μl BV421 anti-CD107a mAb for 2 hr at 37°C (H4A3, Biolegend 328626). NK cells were then stained with mAbs to identify subsets based on the expression of KIR2DL1/S1 (Beckman Coulter, EB6), Receptor (R; Beckman Coulter, Z199, GL183, CD158e1/2) and KIR2DS4 (S4; Miltenyi Biotec, JJC11.6). For intracellular cytokine staining, NK cells were mixed with targets for 6hr at 37°C, Golgi Plug (BD, 555029) was added after 1 hr. Cells were then fixed and permeabilized with Cytofix/Cytoperm (BD, 554714) and stained for IFN-γ (BD, B27) and TNF-*α* (BD, 554512). Cells were washed with PBS three times and data acquired by flow cytometry. To identify KIR2DS1 negative donors, NK cells were stained with mAbs to KIR2DL1 (R&D, 143211) and KIR2DL1/S1 (Beckman Coulter, EB6) as described (Fauriat et al., 2010). For experiments where NK cells were mixed with beads, 10^5^ resting NK cells were mixed with 1.8×10^6^ streptavidin M280 Dynabeads in V bottom 96 well plates, in the presence of 1 μl BV421 anti-CD107a mAb for 2 hr at 37°C (H4A3, Biolegend 328626). Streptavidin M280 Dynabeads were conjugated to biotinylated HLA-C*05:01 refolded with P2-AV or P2-AW as described below.

### Quantification of HLA-C conjugation to beads

Biotinylated recombinant HLA-C*05:01 refolded with P2-AV or P2-AW were supplied by the NIH Tetramer Core Facility. Biotinylated HLA-C (2 μg) was conjugated overnight at 4°C with 6×10^6^ streptavidin M280 Dynabeads in 20 μl. Seven fivefold dilutions of HLA-C were also conjugated in the same way. Dynabeads were washed 5 times with PBS and resuspended in 100 μl PBS for use as targets in NK cell degranulation assays or flow cytometry staining. The density of HLA-C per bead was calculated using QIFIKIT calibration beads according to the manufacturer’s instructions (Dako, K0078). Calibration beads contained five populations of beads conjugated to different numbers of mouse immunoglobulin. Calibration beads were stained with F(ab’)_2_ FITC-Conjugated Goat Anti-Mouse immunoglobulins (Dako, F0479) at 1:50 dilution for 1 hr at 4°C. Beads were washed three times with PBS and analyzed by flow cytometry. The mean fluorescence intensity (MFI) for each bead population was correlated with the number of conjugated mouse Immunoglobulins, provided by the manufacture (Lot 20050787) to generate a standard curve. 10 μl (6×10^5^) of HLA-C conjugated Dynabeads were stained with anti-HLA-I mAb W6/32 (1 mg/ml, Acites) for 1 hr, then washed with PBS three times. The Dynabeads were then stained with F(ab’)_2_ FITC-Conjugated Goat Anti-Mouse Immunoglobulins (Dako, F0479) at 1:50 dilution for 1 hr, both at 4°C. Beads were washed three times with PBS and analyzed by flow cytometry. Using the standard curve generated by the calibration beads, the MFI of W6/32 binding to HLA-C conjugated Dynabeads was used to determine the number of HLA-C molecules per bead.

### Redirected antibody mediated degranulation assays

P815 cells were incubated with the following mAbs alone or in combinations at 10 μg/ml for 20 mins at RT; IgG (MOPC-21, BD Biosciences, 554121), anti-hNKp46 (9-E2, BD Biosciences, 557911), anti-h2B4 (C1.7, Beckman Coulter, IM1607), anti-hCD16 (3G8, BD Pharminogen, 555403) and anti-hKIR2DS4 (179317, R&D systems, MAB1847). Antibody coated cells were mixed with resting NK cells in the presence of 1 μl BV421 anti-CD107a mAb for 2 hr at 37°C (H4A3, Biolegend 328626). Cells were then stained with mAbs to KIR2DS4 (Miltenyi Biotec, JJC11.6, 130-099-963) and KIR2DL1/S1 (EB6, Beckman Coulter, A22332).

### Retroviral transduction of 221 cells with plasmids encoding peptides P2-AV or P2-AW

The peptides P2-AV and P2-AW were expressed in 221–C*05:01 cells using the PresentER system as described (Gejman et al., 2018). The PresentER plasmid encoding a signal peptide from Mouse Mammary Tumor Virus envelope protein followed by the SIINFEKL epitope followed by mCherry was obtained from Addgene (#102945),a kind gift of D. Scheinberg, Memorial Sloan-Kettering, New York, USA (Gejman et al., 2018). The SIINFEKL epitope is encoded by a DNA cassette flanked by mutually exclusive restriction sites for the enzyme SfiI. DNA oligos encoding P2-AV (GGCCGTATTGGCCCCGCCACCTGTGAGCGGGATCATCGACAAGTCCGGCGCCGTGGTGTAAGGCCAAACAGGCC) or P2-AW (GGCCGTATTGGCCCCGCCACCTGTGAGCGGGATCATCGACAAGTCCGGCGCCTGGGTGTAAGGCCAAACAGGCC) with 5’ and 3’ mutually exclusive restriction sites for the enzyme SfiI were synthesized by IGDT. Oligos were amplified using PresentER forward (CGACTCACTATAGGGCCGTATTGGCC) and PresentER reverse (AGTGATTTCCGGCCTGTTTGGCC) primers and cloned into plasmid #102945 following SfiI digestion. 293T Phoenix amphoteric cells in 100 mm plates were transfected with PresentER-P2-AV or PresentER-P2-AW. Virus was collected at 48hr and concentrated with Peg-IT (System Bio) for 48hr at 4°C. 3×10^6^ 221–C*05:01 cells were spinoculated with virus in 6-well plates at 1000g at 32°C for 2hr with 4 μg/ml polybrene. Two days later, transduced cells were selected with puromycin at 0.5 μg/ml.

### Recombinase A protein sequence analysis

Recombinase A protein sequences were downloaded from Genbank. Sequences were searched with the motifs xxD/ExxGxWp9 (x = any residue, p9 = F, L, I, M, V). Duplicate species were removed from returned sequences and the number of unique species and genera were enumerated, for each motif.

### KIR and HLA gene frequencies

Allele frequencies of *HLA-C*05:01*, *HLA-C*04:01* and *HLA-C*16:01* and the frequency of individuals carrying full-length KIR2DS4 (*KIR2DS4*001*) and the KIR2DS4-deleted allele (*KIR2DS4*003*) were obtained from the allele frequencies database (Robinson et al., 2015) (http://www.allelefrequencies.net/). The KIR allele frequencies from 9 populations with high resolution KIR sequencing were from the allele frequencies database (http://www.allelefrequencies.net/), published studies (Gendzekhadze et al., 2009; Nemat-Gorgani et al., 2014; Nemat-Gorgani et al., 2018; Norman et al., 2013; Vierra-Green et al., 2012; Yawata et al., 2006), and kindly provided by Paul Norman (University of Colorado, Denver, USA).

### Analysis of peptides eluted from HLA-C

A total of 2036 9mer peptides eluted from HLA-C*05:01 collected from three studies were used to analyze the frequency of amino acids at peptide position 8 of 9mers, (Di Marco et al., 2017; Kaur et al., 2017; Sim et al., 2017). To compare the frequency of Tryptophan usage in 9mer peptides eluted from 14 HLA-C allotypes, sequences from one study was used (Di Marco et al., 2017).

### Flow Cytometry

Data were acquired on a LSRII or X20 Fortessa, BD. Data were exported as FCS files and analyzed using FlowJo software (Treestar, version 10). Compensation for multicolor experiments were set using single mAb stained beads and Cytometer Setup and Tracking beads were run daily.

### Human donors

Peripheral blood samples from healthy U.S. adults were obtained from the National Institutes of Health (NIH) Department of Transfusion Medicine under an NIH Institutional Review Board-approved protocol (99-CC-0168) with informed consent.

### Statistical analysis

All statistical analysis was carried out in GraphPad PRISM (version 5.0).

## Acknowledgements

We thank P. Norman (U. of Colorado, Denver USA) for HLA and KIR allele frequencies, and the NIH Tetramer Core Facility for recombinant HLA-C. This work was supported by the Intramural Research Program of the National Institutes of Health, National Institute of Allergy and Infectious Diseases, and by a National Institutes of Health–National Institute of Allergy and Infectious Diseases Large Scale Epitope Discovery Program grant (Contract HHSN27220090046C). The funders had no role in the design of the study, data collection, analysis and interpretation, and preparation of the manuscript.

## Author contributions

Conceptualization: M.J.W.S. and E.O.L.; Methodology/Design: M.J.W.S., S.R., P.D.S., and E.O.L.; Investigation: M.J.W.S.; Resources: D.M.A. and R.J.B.; Writing – Original Draft: M.J.W.S., and E.O.L.; Writing – Review & Editing: M.J.W.S., S.R., D.M.A., R.J.B., P.D.S., and E.O.L.; Supervision: P.D.S., and E.O.L.; Funding Acquisition: D.M.A., R.J.B., P.D.S., and E.O.L.

## Additional information

Supplementary Figures 1–7 and Supplementary Table 1 are available online.

---

**Supplementary Figure 1.**
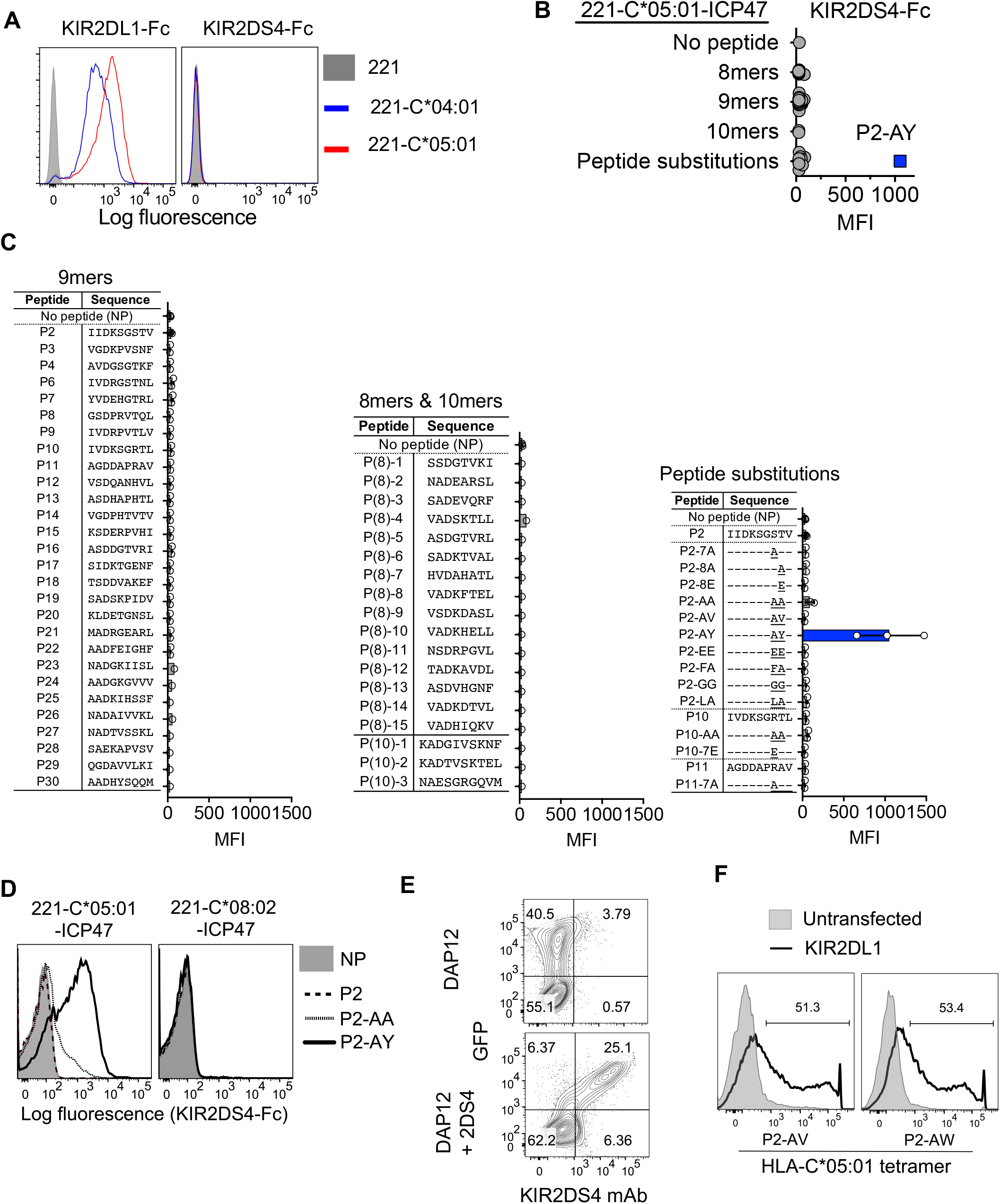
Discovery of a peptide:HLA-C*05:01 complex binding to KIR2DS4. **(A)** KIR2DL1-Fc and KIR2DS4-Fc binding to 221, 221–C*04:01 and 221–C*05:01 cells. **(B, C)** KIR2DS4-Fc binding to 221–C*05:01-ICP47 cells loaded with no peptide (NP) or 28 9mer peptides, 15 8mer peptides, 3 10mer peptides and 13 9mer peptides with amino acid substitutions at positions 7 & 8. Data are summarized in **(B)** and peptide sequences are shown in **(C)**. **(D)** KIR2DS4-Fc binding to 221–C*05:01–ICP47 and 221–C*08:02–ICP47cells loaded with P2, P2-AA and P2-AY or NP. **(E)** Flow cytometry bi-plots displaying KIR2DS4 mAb binding to 293T cells transfected with separate vectors (pIRES2-eGFP) containing cDNA encoding DAP12 and KIR2DS4, or DAP12 only. Binding of anti-KIR2DS4 mAb to 293T cells transfected with DAP12 and KIR2DS4 cDNA or DAP12 cDNA only. **(F)** Binding of HLA-C*05:01 tetramers refolded with P2-AV and P2-AW to untransfected 293T cells or 293T cells transfected with KIR2DL1.

**Supplementary Figure 2.**
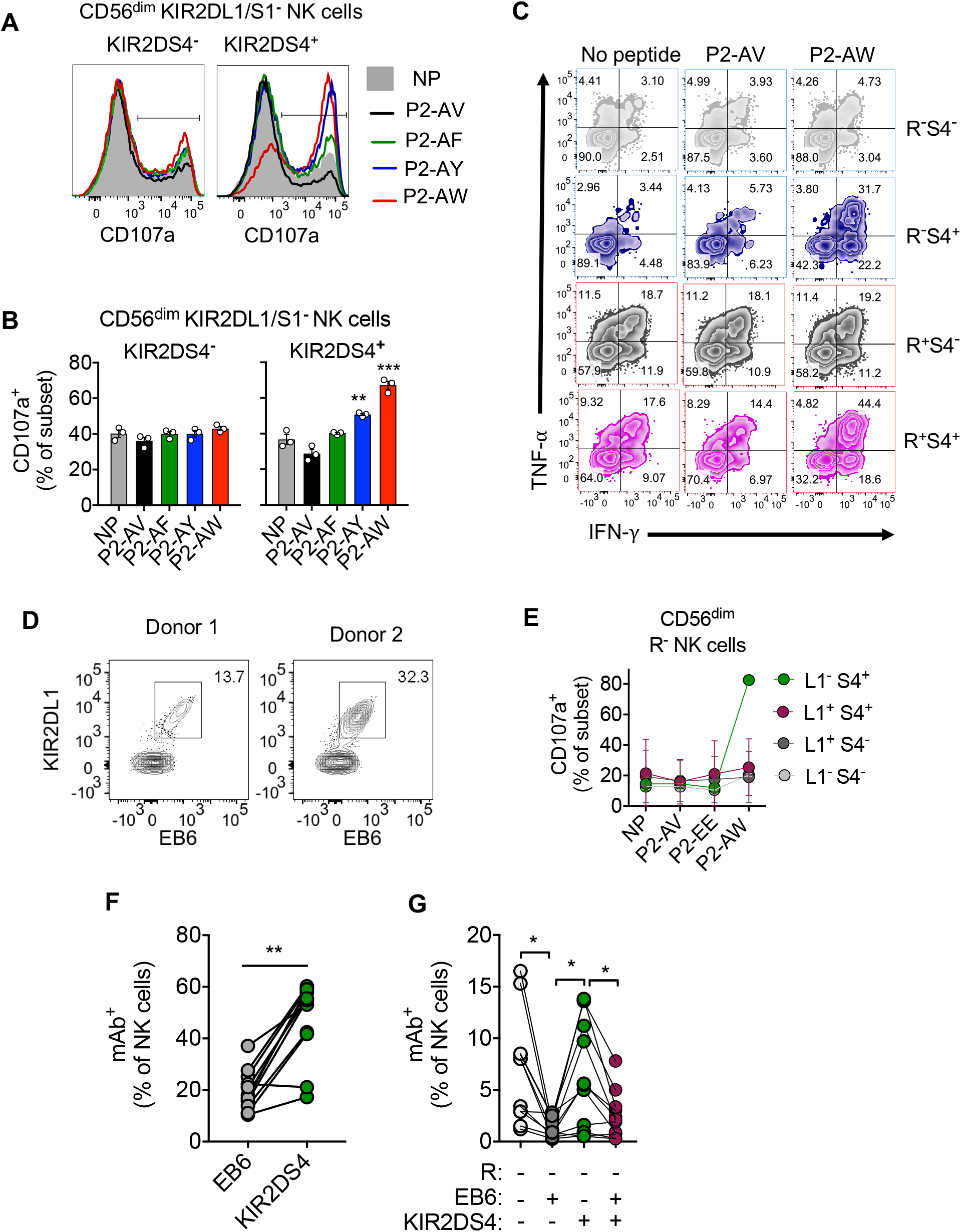
Stimulation of KIR2DS4^+^ cells by peptide:HLA-C*05:01 complexes. **(A, B)** Expression of CD107a on CD56^dim^ KIR2DL1^−^ NK cells in response to 221–C*05:01-ICP47 cells loaded with P2-AV, P2-AF, P2-AY and P2-AW or no peptide (NP). NK cells are gated as CD56^dim^ KIR2DL1/S1^−^. **(C)** Flow cytometry bi-plots displaying expression of IFN-γ and TNF-a on NK cell subsets defined in (Fig. 2A) in response to 221–C*05:01–ICP47 cells loaded with P2-AV, P2-AW, no peptide or no target cells. Data are from one representative experiment out of 3 independent experiments. **(D)** Flow cytometry bi-plots displaying staining of CD56^dim^ NK cells from two donors with a KIR2DL1 specific mAb (143211) and a KIR2DL1/S1 (EB6) mAb. **(E)** Expression of CD107a on CD56^dim^ R^−^ NK cells in response to 221–C*05:01–ICP47 cells loaded with P2-AV, P2-EE and P2-AW or no peptide (NP). NK cells were gated as into four subsets based on expression of KIR2DS4 (S4) and KIR2DL1 (L1). R^−^ NK cells are defined by lack of KIR2DL2/L3/S2 (clone GL183), KIR3DL1/3DS1 (clone p70) and NKG2A (clone Z199). **(F)** Proportion of CD56^dim^ NK cells from 11 donors which express KIR2DL1/S1 (EB6) or KIR2DS4. **(G)** Proportion of CD56^dim^ R^−^ NK cells from 11 donors which express either KIR2DL1/S1 (EB6) or KIR2DS4 or both.

**Supplementary Figure 3.**
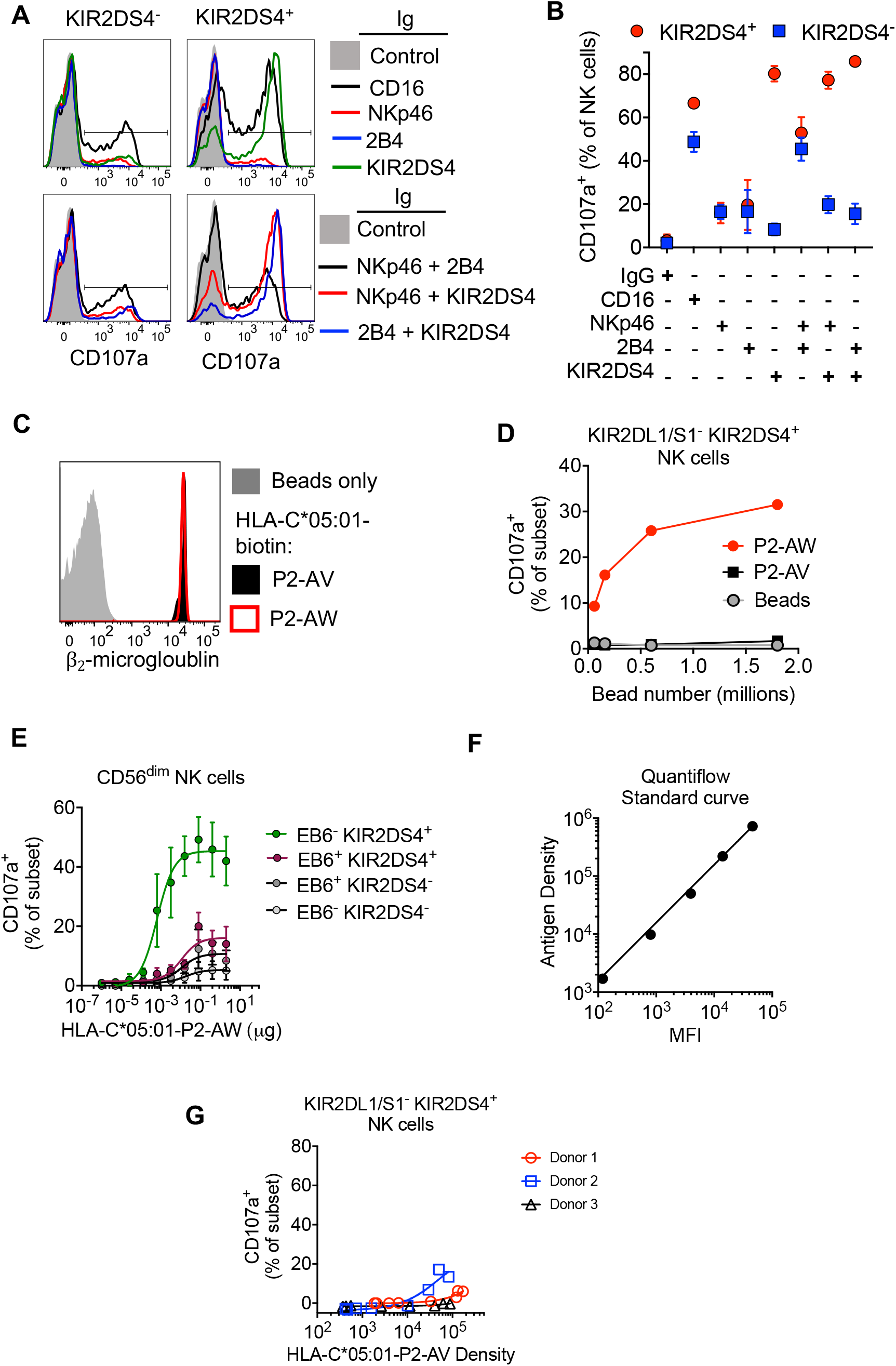
KIR2DS4 is a stronger activator of NK cells than NKp46 or 2B4. **(A, B)** Expression of CD107a on NK cells in response to P815 cells pre-incubated with specific mAbs to CD16, NKp46, 2B4, KIR2DS4 and combinations of mAbs as indicated. NK cells were gated as KIR2DS4^+^ or KIR2DS4^−^ and data from three independent donors are summarized in B. **(C)** Flow cytometry histograms displaying binding of anti-β_2_M mAb to streptavidin Dynabeads only and Dynabeads conjugated to biotinylated HLA-C*05:01 refolded with P2-AV or P2-AW. **(D)** Expression of CD107a on CD56^dim^ KIR2DL1/S1^−^ KIR2DS4^+^ NK cells in response to an increasing number of unconjugated Dynabeads or Dynabeads conjugated to HLA-C*05:01 refolded with P2-AW or P2-AV. **(E)** Expression of CD107a on NK cells in response to Dynabeads conjugated to HLA-C*05:01 refolded with P2-AW. CD56^dim^ NK cells were gated into four subsets based on the expression of KIR2DS4 and KIR2DL1/S1 (EB6). **(F)** Flow cytometry standard curve to determine antigen density from MFI. Calibration beads conjugated to a defined number of mouse immunoglobulins (antigen density) were stained with FITC conjugated polyclonal goat anti-mouse F(ab’)_2_. **(G)** Expression of CD107a on CD56^dim^ KIR2DL1/S1^−^ KIR2DS4^+^ NK cells from 3 donors after stimulation with Dynabeads bearing increasing antigen densities of biotinylated HLA-C*05:01 refolded with P2-AV.

**Supplementary Figure 4.**
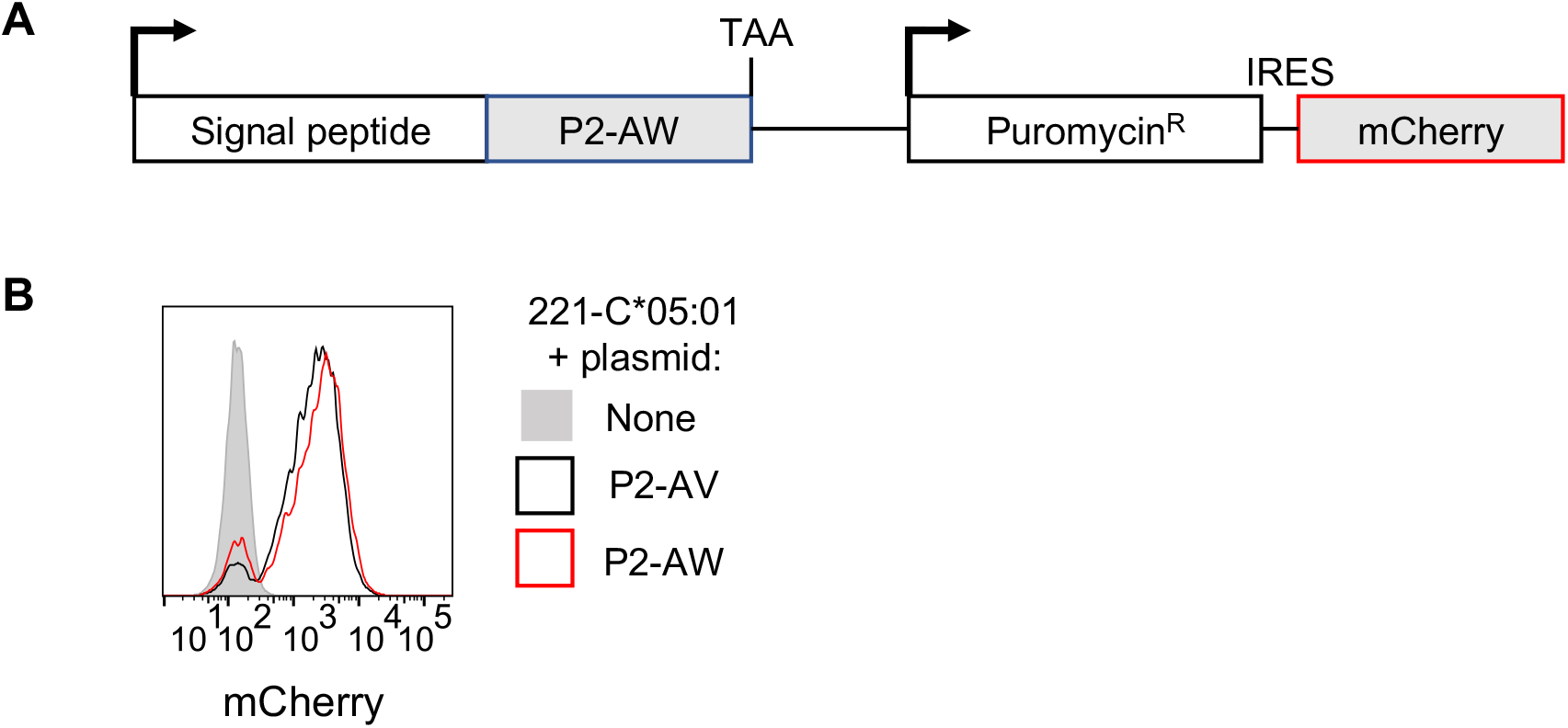
Retroviral transduction of 221–C*05:01 cells with plasmids encoding P2-AV and P2-AW. **(A)** The plasmid encoding P2-AW includes a signal peptide sequence fused to P2-AW. Arrows indicate translation start site, TAA = stop codon, IRES = internal ribosomal entry sequence. **(B)** Expression of mCherry in in 221-C*05:01 cells after transduction with P2-AV and P2-AW expressing plasmids.

**Supplementary Figure 5.**
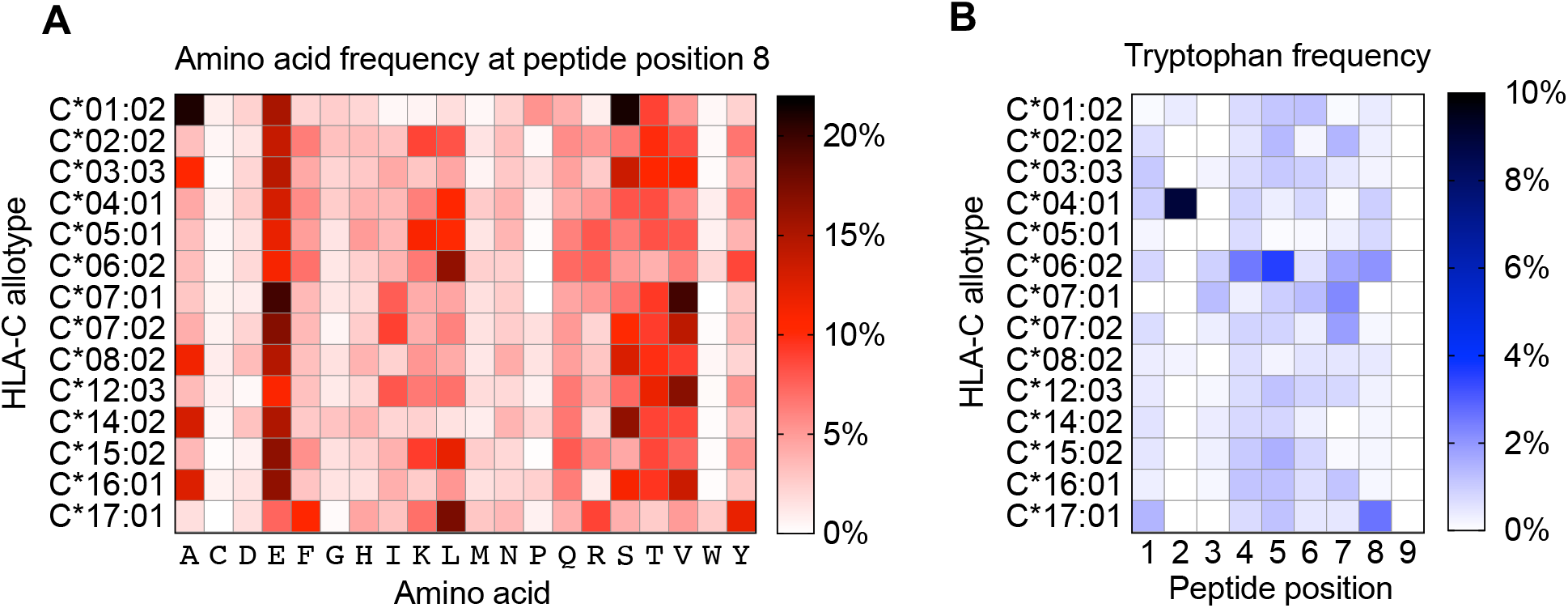
Analysis of peptides eluted from HLA-C. **(A)** Frequency of individual amino acids at position 8 in all 9mer peptides eluted and sequenced from 14 HLA-C allotypes. The number of peptides eluted ranged from 310 (C*07:01) to 1899 (C*16:01). Data from Di Marco et al., 2017. **(B)** Frequency of tryptophan at peptide positions 1 to 9 from all 9mer peptides eluted and sequenced from 14 HLA-C allotypes.

**Supplementary Figure 6.**
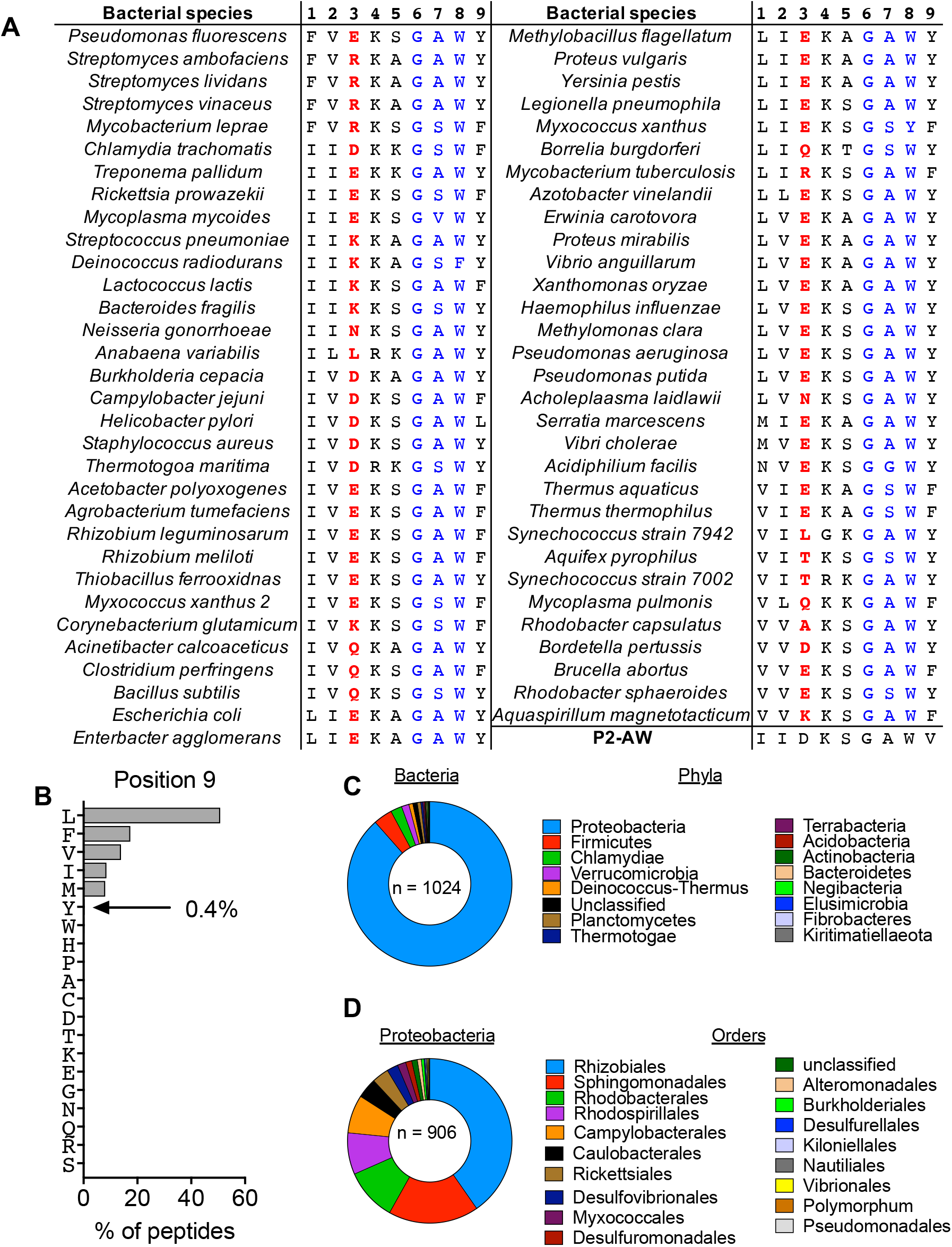
A conserved sequence in RecA contains KIR2DS4 binding peptides. **(A)** Individual RecA_283-291_ sequences from 63 species of bacteria aligned in a previous study (Karlin and Brocchieri, 1996). The highly conserved region at position 6-8 is colored in blue and position 3, essential for binding to HLA-C*05:01 is colored in red. The P2-AW sequence is included for comparison. **(B)** Frequency of individual amino acids at position 9 in peptides eluted and sequenced from HLA-C*05:01. **(C)** Relative abundance of bacterial species (n=1024) in each of the 16 Phyla, which carry a predicted epitope in RecA for binding to HLA-C*05:01 and KIR2DS4. **(D)** Relative abundance of Proteobacteria species (n=906) in each of the 19 Orders, which carry a predicted epitope in RecA for binding to HLA-C*05:01 and KIR2DS4.

**Supplementary Figure 7.**
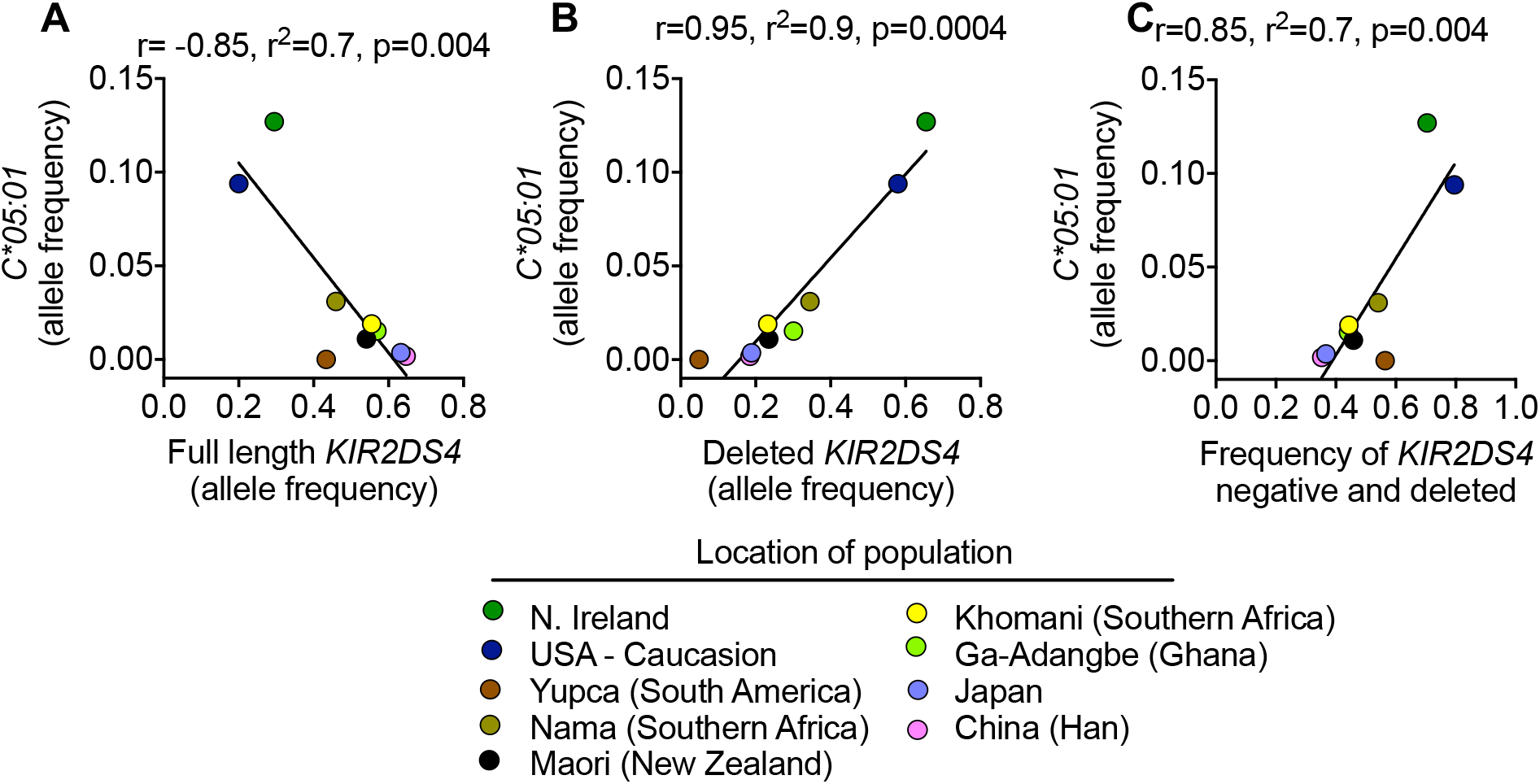
Allele frequency of functional KIR2DS4 is negatively correlated with its ligand HLA-C*05:01. The allele frequency of HLA-C*05:01 in 9 populations is correlated with the frequency of three KIR2DS4 genotypes; KIR2DS4-fl **(A)**, KIR2DS4-del **(B)** and the sum of KIR2DS4-del and those with no KIR2DS4 gene (KIR2DS4-neg; **C**).

